# Citizen science gamers enable automated flow cytometry gating through machine learning

**DOI:** 10.1101/2025.10.07.679685

**Authors:** Sebastiano Montante, Daniel Yokosawa, Leon Li, Alexander Butyaev, Mehrnoush Malek, Razzi Movassaghi, Quentin Michalchuk, Chieh-Ting Jimmy Hsu, David Shmil, Albina Rahim, Andrea Cossarizza, Julia Boira Esteban, Kornél Erhart, Bergur Finnbogason, George Kelion, Hjalti Leifsson, Josh Rivers, David Ecker, Attila Szantner, Jérôme Waldispühl, Ryan R. Brinkman

## Abstract

Manual flow cytometry gating requires up to one hour per sample with 32% inter-expert variability, creating critical bottlenecks in immunological research reproducibility. To address this, we developed flowMagic, a machine learning algorithm for automated gating that is trained on both expert-curated data (template data) and crowdsourced annotations from citizen science gaming. Through *EVE Online*, 839,199 players analyzed 52,178 bivariate plots from 37 studies, generating 31,703 quality-controlled training plots. Evaluated against 92,203 expert-validated files spanning 79 immune populations (i.e., a biologically defined cell subset within each bivariate plot), flowMagic achieved 90% accuracy for abundant populations and 65% for rare populations, outperforming existing methods. The algorithm reproduced biological patterns including neutrophil dynamics in COVID-19 patients and immune development in newborns. This gaming-based approach demonstrates that crowd-sourced pattern recognition generates robust training data for complex biomedical applications, offering transformative potential for standardizing flow cytometry analysis and accelerating immunological discovery.

## Main

Flow cytometry (FCM) is a cornerstone technology in immunology and cell biology, enabling precise identification and quantification of heterogeneous cell populations across diverse applications. Modern immunology increasingly depends on high-dimensional flow cytometry (i.e., flow cytometry experiments involving a high number of biomarkers) to dissect complex immune responses, yet manual gating creates reproducibility crises that undermine translational research. User-defined boundaries are sequentially applied to isolate specific cell populations based on their physical and fluorescent properties^1^. This inherently subjective and time-intensive process yields significant inconsistencies, with a comprehensive evaluation of expert operators demonstrating an interparticipant range in absolute cell population of 32%^2^. Moreover, analysis of a single clinical trial sample can consume up to an hour^3,4^. These inherent limitations become even more pronounced as dataset sizes grow. The scalability constraints of manual gating are increasingly apparent in large-scale clinical trials and high-throughput experiments^1,5–8^, creating an analytical bottleneck that, combined with inherent subjective variability, can profoundly impact research outcomes, clinical decision-making, and scientific reproducibility. These risks include overlooking rare cell populations or subtle marker expression changes, potentially obscuring important biological insights, delaying diagnosis, or inconsistently stratifying patients in clinical settings^6,8^.

Researchers have developed numerous automated algorithms to accelerate and standardize the gating process, addressing the challenges of manual analysis^5–11^. These computational approaches generally fall into two categories: supervised methods, like ElastiGate^11^ (which uses pre-labeled training data) and flowDensity^9^ (which uses panel-specific density-based thresholds), and unsupervised methods such as FlowSOM^12^, which identify cell populations without prior knowledge. Recent efforts have focused on machine learning (ML) techniques, training algorithms with reference examples from specific panels to analyze similar datasets^13,14^. These ML approaches, employing techniques like neural networks, support vector machines and random forests, have shown promise in specific applications such as mature B-cell neoplasm subtyping and acute leukemia classification^13–17^.

However, the development of robust ML algorithms for automated flow cytometry analysis has been significantly hindered by the scarcity of large, diverse and high-quality labeled datasets. As a result these methods often struggle to generalize across diverse flow cytometry panels, typically requiring either complex parameterization or panel-specific examples to achieve optimal results^5,9,18^. While repositories such as FlowRepository^19^ and ImmPort^20^ provide raw data and processed results, there is a critical shortage of large, diverse and high-quality labeled (i.e., gated) datasets essential for training robust and generalizable ML algorithms^21^.

To overcome this data shortage, we leveraged citizen science, an approach used to engage the general public in scientific research^22–24^. We employed an original approach in citizen science to mobilize large crowds of contributors in video games, a concept originally proposed by Massively Multiplayer Online Science (MMOS)^25,26^. Citizen science projects, especially online gaming initiatives, have successfully generated large datasets for ML algorithm training across diverse scientific fields^27–33^. We integrated a flow cytometry gating task into Project Discovery, a citizen science minigame framework existing since 2016 inside the massively multiplayer online game “*EVE Online*”. This platform allows researchers to tap into a massive community of active players, who provide long-term sustained contributions in data analysis. This strategy allowed us to accumulate a vast and varied collection of gated flow cytometry data.The availability of this comprehensive, diverse dataset finally enabled us to overcome the longstanding data bottleneck that has historically limited the development of robust, generalizable automated gating algorithms. With this foundation, we developed “flowMagic”, a novel automated gating algorithm that addresses the field’s core computational challenges.The flowMagic algorithm, trained on a large, diverse set of gated flow cytometry data from multiple research projects, represents a significant advancement in ML-based analysis. This fusion of citizen science and ML addresses the field’s data shortage while enabling more robust, generalizable automated analysis tools.

We developed a rigorous, multifaceted framework for comprehensively evaluating automated gating algorithms in flow cytometry analysis. This framework expands traditional metrics, offering a nuanced evaluation of gating performance across diverse datasets, conditions and cell types. We emphasized challenging low-abundance cell populations by stratifying performance based on population size. This approach assessed the algorithm’s effectiveness across the full spectrum of cell abundance. We also evaluated the reproducibility of biological patterns, a critical aspect often overlooked in quantitative assessments. In addition, we developed expert-validated benchmarks for F_1_ scores, offering a standardized performance measure that balances precision and recall across diverse cytometry datasets. To rigorously evaluate robustness and generalizability, we performed comprehensive assessments across diverse experimental conditions and cell populations, mirroring the complexity encountered in real-world flow cytometry applications. Our evaluation framework addresses key limitations in existing methodologies, establishing rigorous standards for benchmarking automated flow cytometry analysis tools.

## Results

### Citizen Science Data Generation and Quality Control

Our approach leveraged citizen science through the Project Discovery mini-game in *EVE Online* to generate a large dataset of manually analyzed flow cytometry plots (Figure 1). We collected 1,895,367 submissions from 839,199 de-identified player accounts, with 58% related to COVID-19 cases. The dataset underwent rigorous quality control measures, including bot detection, label consistency checks and consensus generation (Supplementary Figure 1-2). This process resulted in a high-quality dataset of 31,703 files, which we then used to train the generalized flowMagic model.

**Figure 1:**
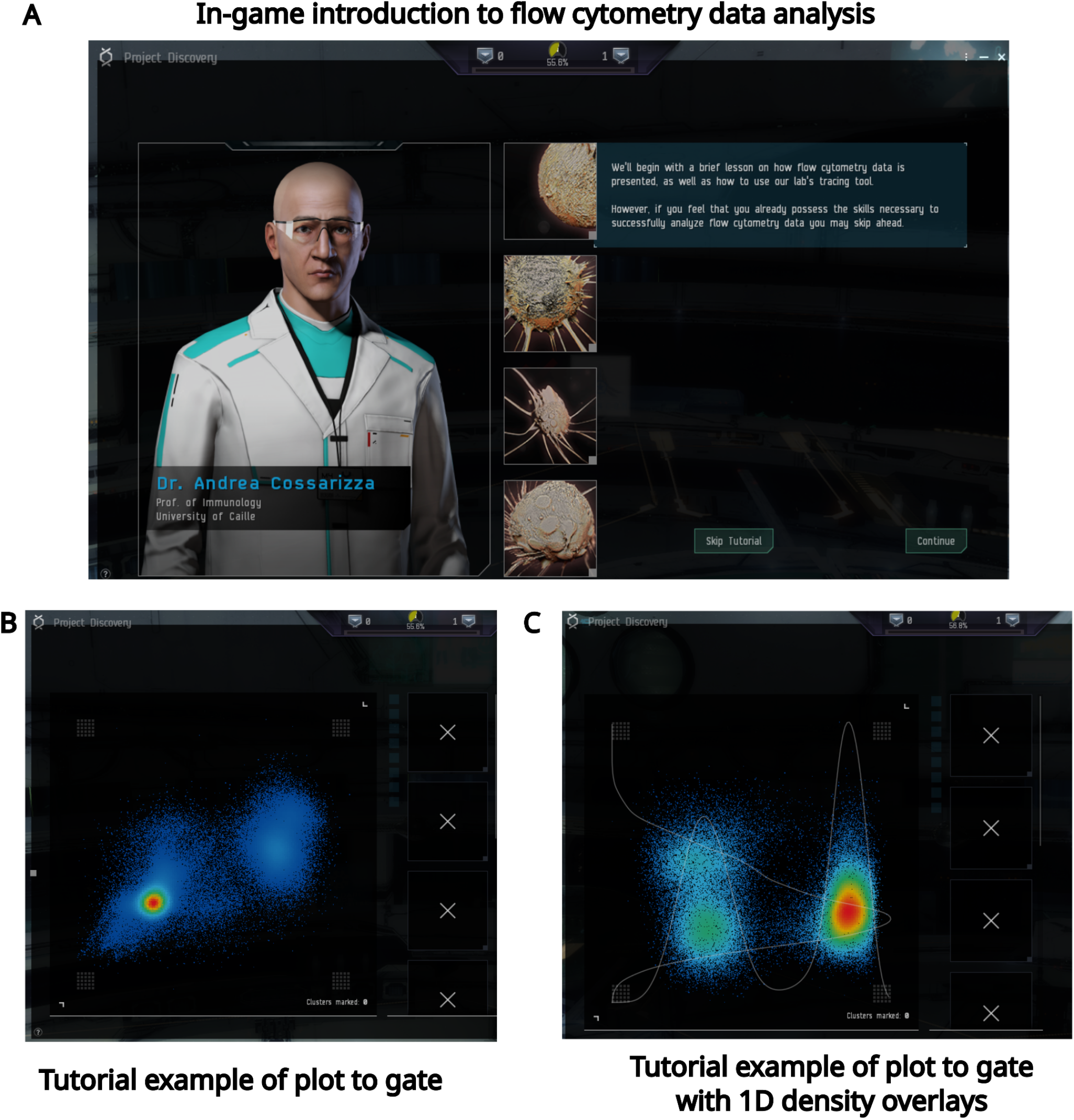
Project Discovery gating mini-game in *EVE online with density overlays*. (A) Introduction page to the Project Discovery gating mini-game with Dr. Andrea Cossarizza (left image) explaining to the players (text on the right) the main concepts of flow cytometry technology and gating. (B) Example of bivariate plot to gate presented to the players during the mini-game tutorial. (C) Example of bivariate plot to gate with one-dimensional (1D) density overlays presented to the players during the mini-game tutorial.

### flowMagic Model Performance

We developed and evaluated two flowMagic models, each one trained only on a specific type of training data: a template model that needs to be trained on expert-gated reference data and a generalized model trained on our citizen science-generated dataset. Our analysis stratified cell populations into large (>2,367 events) and low-abundance (<2,367 events) categories to assess performance across different population sizes. Using flowDensity gates from peer-reviewed studies as the gold standard for F_1_ tests, which removed the subjectivity of human-set gates, the template model outperformed the generalized model, with macro F_1_ scores of 90% vs 78% for large clinically relevant populations (such as T cells and B cells) and 65% vs 42% for low-abundance clinically relevant populations, such as the plasmacytoid dendritic cells (pDCs) (Supplementary Figure 3-8). Both struggled with low-abundance populations and complex, multi-peak gates. Analysis revealed positive correlations between population size and F_1_ scores (Figure 2), and between reference and flowMagic counts (Supplementary Figure 9). A negative correlation between target-template distance and F_1_ scores highlighted the template model’s sensitivity to training data similarity (Supplementary Figure 10-12, Supplementary Note 1). These findings underscore both models’ improved performance with larger populations and the template model’s dependence on relevant training examples.

**Figure 2:**
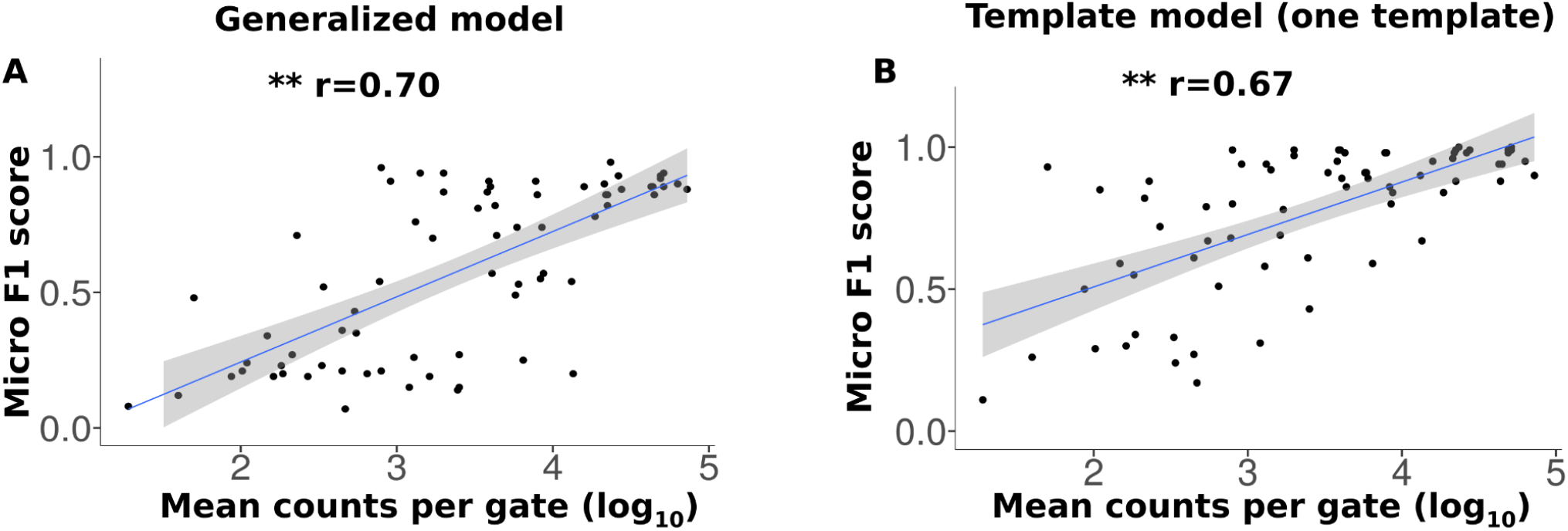
Positive correlation between mean counts and F1 scores. (A) Correlation plot between the micro F_1_ scores generated by the generalized model and the log_10_ of the mean counts per gate. (B) Correlation plot between the micro F_1_ scores generated by the template model trained on one random template and the log_10_ of the mean counts per gate. The grey area indicates the 95% confidence interval of regression predictions. ** p ≤ 0.01 by Pearson correlation test. “r” indicates the coefficient of the Pearson correlation test.

FlowMagic was also compared to the unsupervised gating tool FlowSOM (Supplementary Figure 13-15, Supplementary Table 1) and the supervised gating tool Elastigate (Supplementary Figure 3-4 and 7-8, Supplementary Table 2). flowMagic demonstrated superior clinical performance than Elastigate in identifying very rare pDCs, involved in antiviral immunity and autoimmune diseases (F1 score > 80%), while maintaining accuracy for abundant immune populations essential for routine diagnostics such as B cells (F1 score > 80%). flowMagic also showed superior performance than FlowSOM in identifying T cells (F1 score > 90%), an abundant blood population used to monitor immunoregulation mechanisms.

### Novel Evaluation Approaches

The F1 score is the most common measure for evaluating the performance of automated gating tools. However, there is not a universally accepted consensus about what represents a “good” F1 score. Thus, we used a manual expert-guided approach to estimate an ideal range of F1 scores. We manually assessed 600 random plots across 20 F1 score ranges. Three expert evaluators classified plots as “good” or “bad”, comparing the automated gates with the corresponding reference gates generated and validated by experts in previous studies and evaluating the ability of the algorithms in capturing the previously identified major density regions. Both models showed a consistent positive correlation between F1 scores and the percentage of “good” plots (Figure 3). This trend provides a more interpretable benchmark for F1 scores in flow cytometry analysis, bridging quantitative metrics with expert visual assessment.

**Figure 3:**
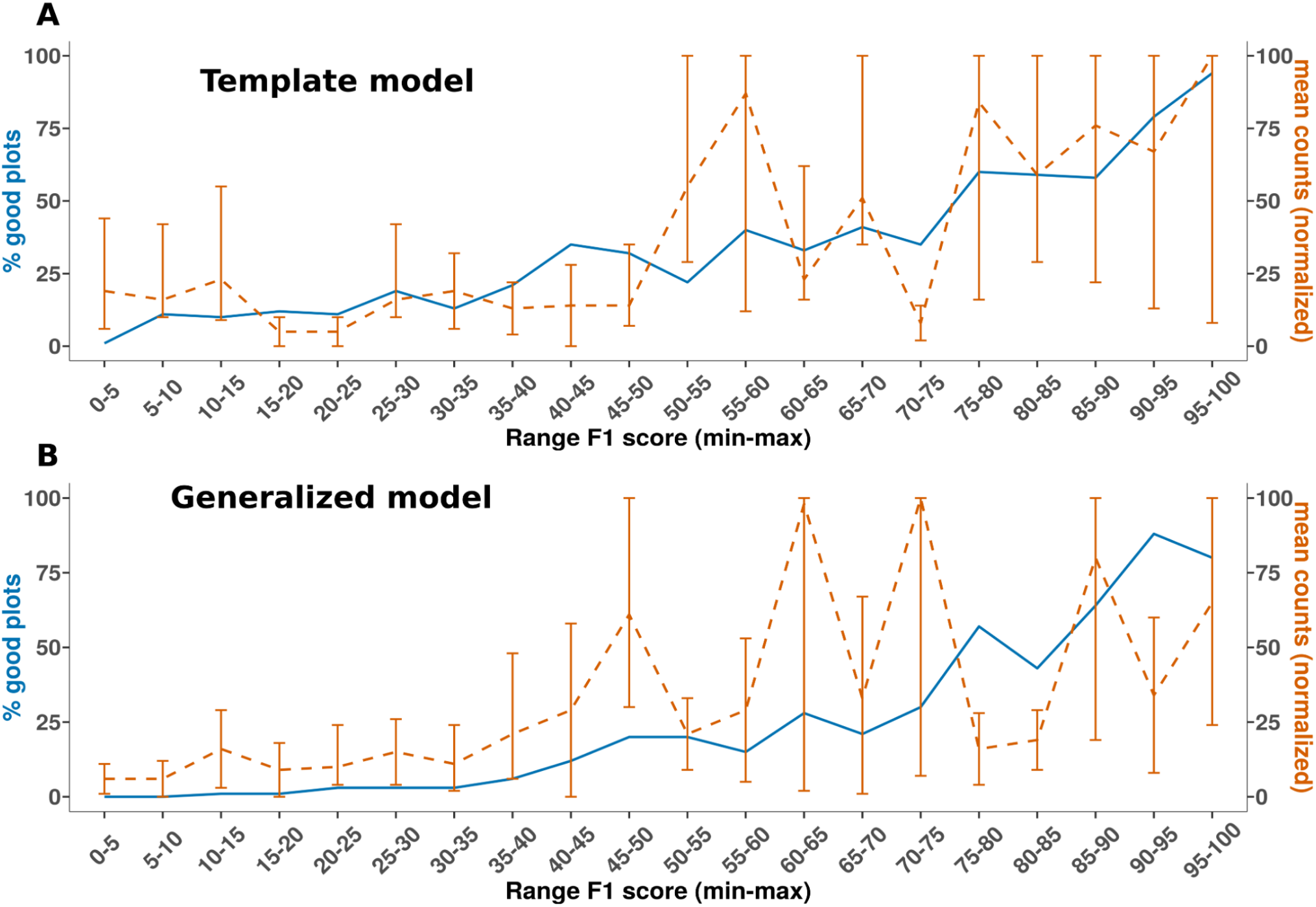
Percentage of good plots across F_1_ score ranges. (A) Template model (trained on one random template) and (B) generalized model (trained on Project Discovery dataset). Blue solid line (left y-axis): Mean percentage of good plots across all researchers for 20 F_1_ score ranges. Orange dashed line (right y-axis): Mean counts of reference gates normalized to maximum count with standard deviation bars. x-axis: F_1_ score ranges, each spanning from the minimum (exclusive) to maximum (inclusive) value for that range.

We evaluated flowMagic’s ability to reproduce biological patterns from previous studies (Figure 4-5). Both models detected key trends in newborn immune development (Figure 4) and identified significant neutrophil increases in non-COVID-19 respiratory infections (p<0.05) in COVID-19 data (Figure 5). However, they missed a marginal increase in COVID-19 patients that was barely significant in the original study (p=0.044).

**Figure 4:**
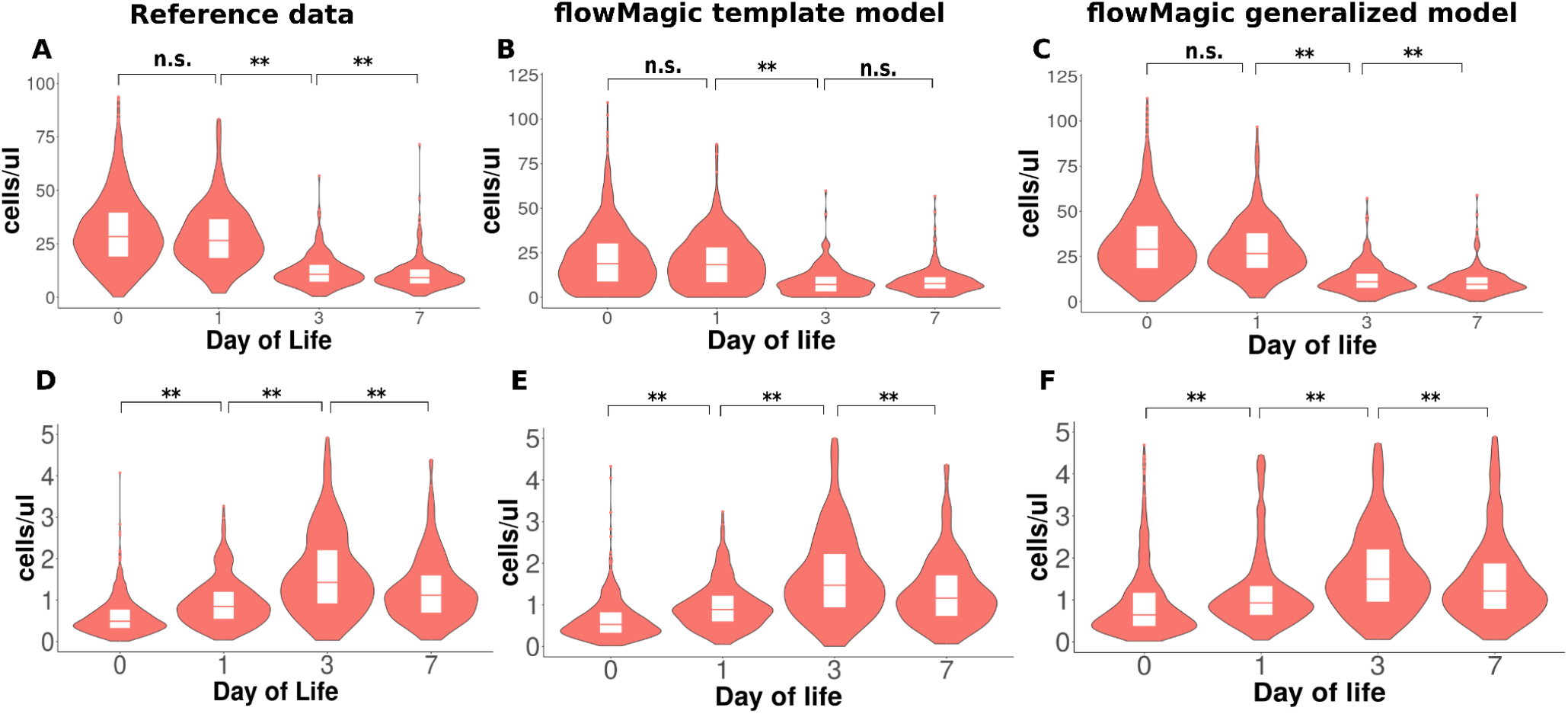
flowMagic identifies biological patterns in FCM data from human newborns. p-values and gated data distribution for (A-C) mature neutrophils and (D-F) mDC; (A,D) Reference data; (B, E) Template model (one template); (C, F) Generalized model; ** p ≤ 0.01 and n.s. p > 0.05 by Wilcoxon Rank-Sum test.

**Figure 5:**
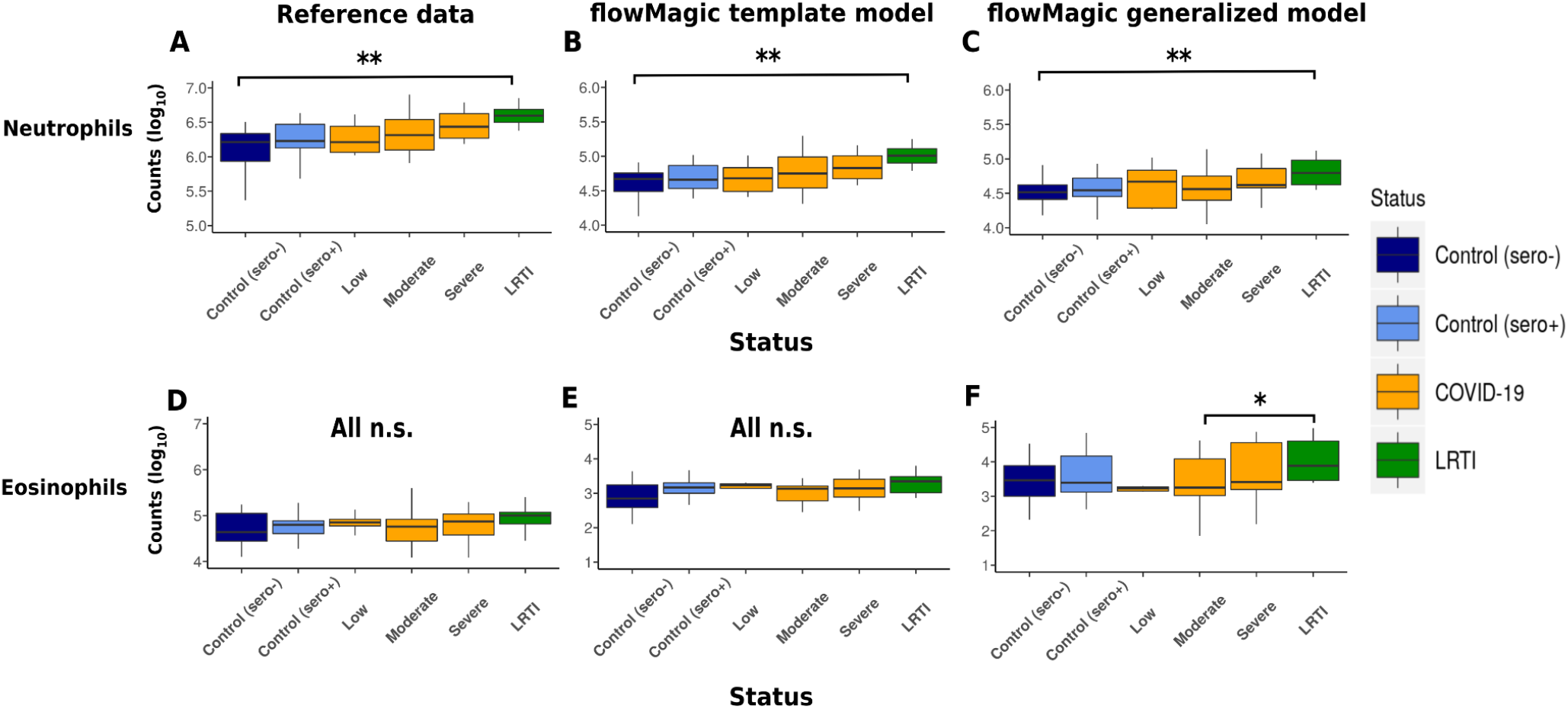
flowMagic identifies reference biological patterns in flow cytometry (FCM) data associated with COVID-19 patients. Reference p-values and gated data distribution for (A) neutrophils and (D) eosinophils; Template model (one template) p-values and gated data distribution for (B) neutrophils and (E) eosinophils; generalized model p-values and gated data distribution for (C) neutrophils and (F) eosinophils. ** p ≤ 0.01 and n.s. p > 0.05 by Wilcoxon Rank-Sum test. Patients grouping include: healthy seronegative patients (Control sero -, blue), healthy seropositive patients (Control sero +, light blue), patients diagnosed with COVID-19 (orange) and patients diagnosed with non-COVID-19 lower respiratory tract infections (LRTI, green).

### Computational Requirements

ElastiGate demonstrated superior speed, gating each plot in <1 second. flowMagic’s template model and FlowSOM required ∼4 seconds per plot, while the generalized model needed ∼6 seconds. Both flowMagic models used 1-2 GB for gating, but training memory requirements differed significantly: 0.5-1 GB for the template model versus 30-100 GB for the generalized model, reflecting its larger training dataset (Supplementary Note 2). FlowSOM required less than 1 GB of memory, while ElastiGate required 10 GB of memory.

## Discussion

Our study introduces flowMagic, a novel machine learning autogating approach that harnesses the power of citizen science to automate flow cytometry gating. By leveraging the pattern recognition capabilities of over 800,000 citizen scientists, we have created a generalized model capable of analyzing diverse flow cytometry datasets with remarkable accuracy. This innovative framework not only challenges the conventional reliance on expert manual gating but also demonstrates the potential of crowdsourced data annotation in addressing complex biomedical challenges and beyond. The interdisciplinary nature of this work (combining immunological domain expertise, gaming psychology and machine learning) was essential for success. The challenge of generating high-quality, diverse, scalable and well-curated training datasets extends far beyond flow cytometry, across various high-risk fields including healthcare, cybersecurity and defense^34,17^. Strategies that ensure dataset diversity, accuracy, and robustness are essential for developing reliable AI systems in any domain where model performance critically depends on the quality and variety of labeled data^34,17^.

The use of a gaming platform (specifically through Project Discovery in *EVE Online*) is uniquely suited to the pattern-based nature of flow cytometry gating. Gating often involves recognizing distinct density distributions and visual boundaries within two-dimensional plots, a task that closely resembles visual and spatial challenges commonly found in video games. Players are naturally adept at solving such tasks, especially when embedded in gamified environments that reward pattern recognition, consistency, and quick decision-making. Unlike traditional crowdsourcing methods that may lack sustained engagement, games provide intrinsic motivation and scalability, enabling rapid accumulation of high-volume, diverse, and quality-controlled annotations. Moreover, the integration into an existing online game allowed us to bypass many logistical and engagement barriers that plague conventional citizen science platforms, enabling faster deployment and broader demographic reach.

Pure computational approaches fail to capture biological nuances, while traditional immunology lacks scalable analysis tools. The performance of flowMagic, particularly in identifying large cell populations, rivals that of current state-of-the-art methods while offering increased speed and scalability. flowMagic’s bivariate results are easy to interpret for the majority of scientists who tend to prefer the definition of target populations in the traditional bidimensional approach^2^. Evaluating automated gating solutions against manual gating as the gold standard using the F_1_ score creates a methodological contradiction, as it attempts to validate computational methods that were specifically developed to address human subjectivity and variability against those same subjective human annotations, essentially using the problem to validate its solution. Our key insight is that while absolute ‘truth’ in gating may be subjective, we can systematically engineer controlled perturbations where we precisely know the magnitude of change between states. This allows us to evaluate gating tool performance by measuring how well they track these known changes, rather than how well they match manual gates.

While the flowMagic template model achieved higher overall accuracy (in both high and low-abundance populations), the generalized model demonstrates remarkable versatility. It handles diverse gating scenarios without panel-specific expert templates, effectively reproducing biologically significant patterns in newborn and COVID-19 studies. Unlike template models that may force data into predefined patterns, the generalized model’s flexibility allows adaptation to diverse density distributions. The generalized model displayed superior performance versus ElastiGate on low-abundance populations, an area where existing tools often struggle^8,18,4,35^, suggesting that the diversity and scale of its training data may confer advantages in detecting and accurately gating rare cell types. Rare populations are defined as populations having low frequency of cell counts, usually less than 5% of the parent population^8^, showing higher variance when analyzed by automating gating methods^8^. In our evaluation method, instead of setting an arbitrary threshold, we defined the rare populations based on the empirical distribution of population sizes in our data, estimating a data-driven threshold, rather than an arbitrary cut-off. By setting the threshold at the median of mean population sizes, we effectively separate larger populations and rare populations in a way that is consistent across datasets, without relying on an arbitrary percentage that may vary depending on sample composition.

By capturing a wider range of gating scenarios, the generalized model could uncover biological patterns overlooked by rigid methods. Of note, ElastiGate may provide superior accuracy with proper parametrization (e.g., tuning the density level option in the FlowJo plugin), but our tests aimed to evaluate gating performance without parameter optimization. Furthermore, while both ElastiGate and flowMagic template models were designed to reproduce only the gates present in template data, ElastiGate has an additional limitation: when provided with multiple templates containing different marker expression and gates, Elastigate selects a single training image based on similarity to the plot to be gated and it fits gates using only that example. In contrast, the flowMagic template model incorporates all gates and expression patterns from the complete training dataset.

Both the template and generalized approach struggled with low-abundance populations. A key differentiator of our approach is the explicit consideration of cell population size when evaluating algorithm performance. Unlike previous studies that rely solely on F_1_ scores, we stratified populations into large and low-abundance categories, revealing significant performance differences. This approach acknowledges that population abundance significantly affects identification accuracy and is likely a primary determinant of performance. For instance, both the template and generalized models achieved higher F_1_ scores for larger populations, with notable struggles in identifying low-abundance populations likely due to their higher variability (we discuss this aspect in more detail in Supplementary Note 1, 3 and 4). By incorporating population size into our analysis, we provide a more nuanced and realistic assessment of algorithm performance. This strategy offers a better understanding of each model’s strengths and limitations across different population sizes, a crucial factor often overlooked in previous evaluations using F_1_ scores alone. Our findings underscore the importance of considering population abundance when developing and assessing flow cytometry analysis algorithms, potentially guiding future improvements in the field. The development of specialized techniques for low-abundance population identification, possibly incorporating additional contextual information or leveraging advanced machine learning architectures designed for imbalanced data, could significantly advance the field.

FlowSOM is one of the most used and accurate automated gating tools^4,18,35^ that can work on a multidimensional space (using all markers, as originally intended by the authors). We tested FlowSOM on both a bidimensional and multidimensional space, with the latter showing an inferior macro F1 score (< 50%), suggesting that the bidimensional approach can offer superior accuracy, in addition to better interpretability. Template-based models, specifically flowMagic and ElastiGate, demonstrated superior performance in gating multi-peak populations compared to flowMagic generalized model.

These template models achieved macro F1 scores exceeding 80% due to their training on expert-defined expression patterns. Our findings suggest that automated gating algorithms trained on expert-gated data offer the most effective approach for delineating multi-peak populations in flow cytometry analysis, as widely expected since the generalized model was trained on a citizen science dataset composed of populations determined by single density peaks. In addition, the template model matched FlowSOM’s execution time, while the generalized model was marginally slower, taking an additional minute to process the same number of files.

Despite the expected superiority of the template model with pre-defined populations, these comprehensive results demonstrate the potential of our citizen science approach in generating valuable training data for automated flow cytometry analysis aimed at identifying unknown populations in FCM bivariate plots. The flowMagic template model approach confirmed its expected superior performance with pre-defined populations, additionally overcoming the performance of other recently developed template approaches. However, the template-based approach required manual exploration of the dataset to find suitable templates, while the generalized model enables automated gating without designing specific templates, offering the possibility of a more unbiased approach.

Our results highlight the strengths and limitations of our flowMagic algorithm, particularly its performance across different population sizes and its ability to preserve biological patterns. The novel evaluation methods we introduced provide a more nuanced understanding of gating tool performance, setting a new standard for assessment in the field of flow cytometry analysis.

The scale and diversity of our training dataset addressed limitations of previous automated gating approaches, which often relied on smaller or less diverse training sets. The success of our citizen science approach raises important considerations about data quality and the potential for crowdsourcing in other areas of biomedical research. The rigorous quality control measures we implemented, including bot detection and consensus generation, were crucial in ensuring the reliability of our training data^36–40^. However, the variability in player skill and engagement presents both challenges and opportunities for future iterations of this approach. More sophisticated player rating systems or adaptive task assignment could further optimize data quality and potentially expand the complexity of tasks that can be crowdsourced.

It is crucial to acknowledge the limitations of our approach. The reliance on non-expert participants necessitates careful task design and extensive quality control measures. We recently added 1-dimensional (1D) density overlays to the display and found this significantly increased the pass rate from 1.5% to 4.45% for bivariate plots with 500+ users (p < 2.2e-16) across 192,000 plots analyzed with the new visualization. Due to data privacy agreements, there is no way to further separate bots from poor human analysis. We hypothesize the addition of 1D densities would aid manual gating for cytometrists by guiding placement of gate boundaries in challenging regions. Additionally, while our generalized model showed impressive performance across a wide range of gating scenarios, there may be highly specialized or extremely low-abundance cell populations that require expert knowledge to identify accurately. Future work should explore the boundaries of what can be effectively crowdsourced and develop hybrid approaches that optimally combine citizen science contributions with expert knowledge (Supplementary Note 3-4). The performance and flexibility of the two versions of flowMagic reveals both the strengths and limitations of current automated gating approaches. The generalized model’s performance is likely constrained by its reliance on top-level bivariate pairs, underscoring the challenge of fully capturing the complexity of multi-level gating hierarchies. However, the consistency in bivariate plot structures across different markers and cell types continues to support the promise of model-based automated gating. The superior performance of the template model, attributed to its specificity and alignment with expert knowledge, highlights the value of incorporating domain-specific expertise into automated analysis tools. The performance of the generalized model was significantly limited by the absence of gates composed of multiple density peaks in the citizen science training data, a task that is difficult to teach non-expert users due to its reliance on prior biological knowledge. To prevent major inconsistencies arising from the challenge non-experts face in learning complex patterns, we intentionally excluded the possibility of populations composed of multiple peaks. Our data quality control pipeline admits only consistent gates composed of single peaks, filtering out those with multiple peaks (Supplementary Note 5). The possibility to generate (with the support of expert knowledge) a large training set that includes populations composed of multiple density peaks, would dramatically increase the accuracy and possible applications of the generalized model (Supplementary Note 3).

As we confront the increasing intricacy of flow cytometry analysis, the lack of field-wide consensus on optimal gating strategies and annotation of cell populations presents a significant hurdle. The majority of automated gating tools analyze the multidimensional dataset (all markers at a time)^4,18,35^. The Flow Cytometry: Critical Assessment of Population Identification Methods (FlowCAP) challenges were established to compare the performance of computational methods for identifying cell populations^41^. The results of these challenges showed that the reliance on classically defined cell types for manual data analysis is problematic in high-dimensional experiments, as the definition of a particular cell type might differ by investigator or paper^41^. To address these challenges, we are expanding the Project Discovery citizen science infrastructure to create comprehensive reference maps for flow cytometry data annotation through the open science SOULCAP Initiative. This collaborative effort, uniting leading academic flow cytometry societies, institutions, pharmaceutical companies, reagent manufacturers, and software developers, aims to establish standardized cell population identification and annotation. By combining large-scale citizen science data generation with expert-driven standardization efforts, we anticipate significant advancements in automated flow cytometry analysis, potentially revolutionizing our ability to identify and characterize diverse cell populations across a wide range of applications in immunology and cancer research. Data collection has steadily continued, with over 647-million plots now analyzed by over 1-million accounts across 1,891 days.

## Methods

### Project Discovery Data Generation and Quality Control

We integrated flow cytometry gating tasks into the massively multiplayer online role-playing game “*EVE Online*” as a mini-game called Project Discovery (PD), utilizing the scientific data collection pipeline of a Massively Multiplayer Online Simulation (MMOS). Players earned in-game rewards for completing gating tasks, incentivizing participation in this citizen science initiative. An in-game tutorial provided examples of correctly gated bivariate plots, referred to as “gold standards”, to prepare players for the task (Supplementary Figure 1).

From a comprehensive collection of 69 flow cytometry datasets, we curated a final set of 37 for analysis (Supplementary Table 3). These datasets were sourced from public repositories including flowRepository^19^, ImmPort^20^, as well as from partner researchers. The selection process excluded datasets lacking essential marker information, those without compensation files, and CyTOF-only datasets, ensuring a high-quality and relevant corpus for our study. This rigorous curation process allowed us to focus on datasets that were most suitable for our analysis objectives, balancing diversity with data quality and completeness.

The samples were acquired in FCS format from each public repository. Based on the hierarchy information reported in the WSP files generated by the FlowJo framework^11^, the events associated with the first level of the hierarchy were extracted from each FCS file. The first level analyzed corresponds to the first cell population of the hierarchy after the removal of technical artifacts (e.g., doublets) and dead cells.

To enhance data diversity and improve the quality of the dataset^34,42^, we applied the flowSim tool^43^ to the bivariate data from each dataset, executing multiple cycles until the heterogeneity score exceeded 0.90. The resulting dataset comprised 52,178 bivariate plots distributed to players for gating.

Players submitted 1,895,367 gated plots from 839,199 accounts, with about 58% related to COVID-19 cases. We implemented a three-step quality control process to ensure high-quality data generation. First, we developed a multi-step bot detection process to identify and remove data gated by automated systems (Supplementary Note 5). Next, we performed a label consistency check on the de-botted dataset, ensuring consistent labeling based on gate locations (Supplementary Note 6). Finally, we used the SpectralClustering algorithm from the scikit-learn library^45^ to generate consensus results from multiple player inputs for each plot (Supplementary Note 6).

The final high-quality dataset comprised 31,703 files, serving as the training set for flowMagic’s generalized model. We also retained a larger dataset of 575,396 files without the final consensus step to evaluate the efficiency of consensus in improving prediction accuracy.

### flowMagic Algorithm Development

The flowMagic algorithm, developed as an R package, consists of four main steps: input loading, pre-processing, gating, and post-processing (Supplementary Figure 16-18). It utilizes 36 training features based on event expression and density. After testing several machine learning models, including Random Forest (RF), feed-forward neural network (FNN), K-nearest neighbor (KNN), decision tree (DT), and Naive Bayes (NB)^46^, we selected the Random Forest model for its superior performance (Supplementary Table 4, Supplementary Note 7).

We employed out-of-the-bag cross-validation for the template model and repeated k-fold cross-validation (k=2, 100 repetitions) for the generalized model (Supplementary Note 7). To manage computational resources, we applied downsampling to 500 points for the generalized model. The caret package in R^47,48^ was used for training and prediction steps, optimizing computational time and memory usage (Supplementary Note 7).

### Evaluation of automated analysis approaches

We evaluated flowMagic’s performance using four datasets: the HIPC dataset^48^ (1,379 samples for Innate panel, 1,382 for Adaptive panel), the OneStudy dataset^3^ (30 samples for Basic panel), the International Mouse Phenotyping Consortium (IMPC) dataset (1,280 samples for Panel 1), and COVID-19 patient data^49^ (226 samples). In total, we analyzed 4,297 samples and 80 cell populations across 39 different expression patterns, encompassing 92,203 bivariate plots - the largest scale evaluation of an automated gating tool to date.

We compared flowMagic’s performance to FlowSOM and ElastiGate. FlowSOM is among the most cited unsupervised gating tools showing the highest speed and accuracy^4,18,35^. FlowSOM represented unsupervised gating tools, while ElastiGate represented supervised gating tools integrated within the popular FlowJo framework^11^. We performed the testing of FlowSOM on two markers at a time using the cytocluster Python package^50^ which provides an optimized framework for applying clustering algorithms on FCM data (it optimizes the execution of FlowSOM on bidimensional data). Parameters were set as default (maximum number of clusters = 50). We also tested FlowSOM on multidimensional data (all markers at once) using the FlowSOM R package^12^, setting a predefined number of clusters based on the expected populations as suggested by the package documentation.

Our evaluation employed multiple methods to assess gating accuracy. We calculated both micro and macro F1 scores^51^ to evaluate performance across different population sizes and types. In particular, the cell populations were divided in two types based on the number of events: large populations (> 2,367 events) and low-abundance populations (< 2,367). The threshold for the cell counts was determined as the median of the vector composed of all means for all cell populations. Specifically, we chose this threshold measure in order to create a data-driven and balanced distinction between “rare” and “large” populations. This approach avoids setting an arbitrary cutoff and ensures that the classification reflects the empirical distribution of cell counts across the dataset. By using the median of means, we aim to separate populations into two roughly equal groups, capturing the natural variation in abundance while enabling consistent stratification for performance evaluation. We also assessed flowMagic’s ability to reproduce significant biological patterns identified in previous studies, focusing on newborn data and COVID-19 patient data.

Additionally, to establish a benchmark for “good” F1 scores, three researchers manually evaluated 600 random plots across 20 ranges of F1 scores. This comprehensive evaluation framework aimed to provide a nuanced understanding of flowMagic’s performance across various gating scenarios and population types, setting a new standard for the evaluation of automated gating tools in flow cytometry analysis while respecting current standards in ethical data collection for the training of generalized algorithms (Supplementary Note 8)^52^.

## Data availability

Project Discovery bivariate training plots used to train the ML model is available from https://www.frdr-dfdr.ca/repo/dataset/fe62f923-d7f4-4fd8-9ff1-5bcd9e6f519b under the Creative Commons Attribution 4.0 International License. The flowMagic R package code along with its documentation is available from Github under the Apache 2.0 license: https://github.com/semontante/flowMagic. The flowMagic evaluation results and the generalized model trained on the players data is available from the https://doi.org/10.20383/103.01352. Full methodology is available on Supplementary Material.

### Ethics declarations

This study follows the most recent Human-centric computer vision (HCCV) data curation practices to assure the generation of a high quality unbiased dataset (see Supplementary Note 8 for additional details).

Attila Szantner is the CEO and founder of MMOS, a Swiss innovator company that creates novel citizen science solutions and provides software that enables citizen science within video games, like Borderlands Science in *Borderlands 3* and Project Discovery in *EVE Online*. Julia Boira Esteban, Bergur Finnbogason, George Kelion, Hjalti Leifsson, Josh Rivers, David Ecker are employees of CCP Games. Kornél Erhart is a collaborator of MMOS. CCP Games develops the game *EVE Online* and the citizen science game *Project Discovery. Project Discovery* is a free mini-game available within the *EVE Online* game that does not require an *Eve Online* premium subscription to play.

## Acknowledgements

We thank all the players of *EVE Online* that contributed to the generation of the data used in this study. This study was funded by New Frontiers in Research Fund NFRFR-2021-00300 and NSERC. The development of the MMOS scientific data pipeline was supported by the European Union’s Horizon 2020 research and innovation programme under grant agreement Nr 732703.

## Authors’ Contribution

Sebastiano Montante developed the flowMagic algorithm, performed and organized the evaluation testing, interpreted the results and wrote the manuscript with Ryan Brinkman, who conceived and supervised the project. Daniel Yokosawa performed the preprocessing, postprocessing and collection of the Project Discovery data. Leon Li developed the bots detection and labels consistency steps of the QC pipeline. Razzi Movassaghi developed the final aggregator step of the QC pipeline. Mehrnoush Malek contributed to Project Discovery data preprocessing and collection. Quentin Michalchuk contributed to the execution of the flowSim algorithm. Chieh-Ting Jimmy Hsu contributed to the processing of Project Discovery data. David Shmil contributed to the evaluation testing of flowMagic. Albina Rahim provided the IMPC test data. Andrea Cossarizza provided the initial COVID-19 data. Alexander Butyaev, Julia Boira Esteban, Kornél Erhart, Bergur Finnbogason, George Kelion, Hjalti Leifsson, Josh Rivers, David Ecker, Attila Szantner and Jérôme Waldispühl developed the *EVE Online* project Discovery infrastructure. All authors reviewed and approved the paper for submission.

