## Supplementary_Material for "Citizen science gamers enable automated flow cytometry gating through machine learning"

### Supplementary Note 1: Data Variability and Template Selection

Two key factors influence the number of required templates: data variability and the importance of this variability in relation to gate boundaries.

To address data variability, we propose using the heterogeneity score calculated by flowSim, a tool designed to identify and remove similar FCM bivariate images. Users should extract bivariate data for each gating step and execute flowSim separately on the data related to each step. For example, if 80% of a cell population's bivariate data falls into one group identified by flowSim, a single example may sufficiently cover most of the variability. Conversely, if data is evenly split between two groups, more templates would be necessary to cover the variability adequately.

To select ideal templates, we recommend running flowSim multiple times until the heterogeneity score reaches 0.9.

Regarding variability importance, populations with inconsistent gate boundaries require more templates (Supplementary Figure 11), while a single template often suffices for populations with consistent gate boundaries (Supplementary Figure 12). For inconsistent gate boundaries, density changes directly translate into differences in gate boundaries (Supplementary Figure 11.A and 11.D). In contrast, consistent gate boundaries remain unaltered by density changes (Supplementary Figure 12.A and 12.B).

Further confirming this, for inconsistent gate boundaries, the target-template distance influenced accuracy (lower micro F1 score with higher distance, see Supplementary Figure 11). For consistent gate boundaries, the target-template distance had no influence (consistent micro F1 score regardless of distance, see Supplementary Figure 12). Note that Elastigate showed a low performance on the Effector CD4+ NKT population (bimodal distribution of F1 scores, with a large group showing low performance), indicating that it was more influenced by the irrelevant density variation of its consistent gates. The flowMagic tool trains on both expression and density features. Thus, it is trained to learn when the expression is more important than the variation in the density features and vice-versa.

As demonstrated in the results section, rare populations were more affected by the number of templates. This is likely because most rare populations exhibit inconsistent gates or higher variability.

### Supplementary Note 2: flowMagic Technical performance

**Gating Performance.** Gating speed varied depending on the model type. The template model required ~4 seconds per plot (without downsampling), comparable to the FlowSOM tool. The generalized model (with downsampling to 500 or 2,000 points) required 4-12 seconds per plot, with increased time due to necessary post-processing. Elastigate demonstrated significantly faster gating speeds than flowMagic and FlowSOM, requiring approximately 0.3 seconds per gate in a bivariate plot. In terms of computational resources, memory usage during gating was consistently 1-2 GB, irrespective of the model, using default downsampling parameters.

**Model Training**

Training time varied considerably based on model type, number of plots, and event count. The template model (without downsampling) required 1-2 minutes, while the generalized model (with downsampling) took up to 1 hour. Activating downsampling for the template model reduced training time to seconds. Downsampling is recommended for the template model when training more than 200 templates to optimize execution time and memory usage.

Memory requirements during training were model-dependent. The template model required 0.5-1 GB, while the generalized model necessitated 30-100 GB with downsampling, requiring high-performance computing resources. Training the generalized model without downsampling exceeded available memory limits.

The resulting generalized model size ranged from several MB (for approximately 100 plots trained) to 3 GB (for 9,170 plots trained).

### Supplementary Note 3: Improving Project Discovery dataset

To enhance the Project Discovery (PD) dataset, we propose generating multiple specialized datasets, each focusing on a specific cell population or gating boundary of interest. For instance, separate datasets could be created for identifying plasma cells, non-granulocytes, or technical artifacts like singlets using the scatter channels (like the FSC/SSC channels). This approach would allow for the inclusion of expert-supervised gates that were previously excluded from the non-expert-generated PD dataset.

The plots showing the univariate density of the data for each marker improved player accuracy. Preliminary results showed an increase in accuracy of 1.50% using plots with events with univariate density next to each axis. Future versions of the PD dataset generated by showing plots with univariate density could improve the quality of the training data.

Rather than a single global dataset, multiple global datasets could be trained separately, as the features determining cell populations would be dataset-dependent. This process would require dedicated instructions and quality-checking algorithms for each cell population, demanding more time and resources. However, it would address the current lack of complex boundaries in the dataset.

It is important to note that the current PD dataset was generated using only the first level of the gating hierarchy from each study, typically containing the largest populations. Expanding to lower hierarchy levels could enrich the training set with gates for smaller populations.

This approach would provide a more comprehensive and nuanced dataset, potentially improving the performance of machine learning algorithms in identifying diverse cell populations and gating boundaries.

### Supplementary Note 4: Potential Improvements and Limitations of flowMagic Algorithm

**Expanding feature set:** The flowMagic prediction accuracy could potentially be improved by expanding the feature set. New features might include the length of a tail and its exact starting point, the presence of twin peaks in a plot, or the presence of a shoulder before a peak. A more refined feature engineering process could further increase the accuracy of the flowMagic algorithm, particularly for the generalized model, which faces the challenge of learning a larger variance of patterns compared to the template model.

**Limitations in improving certain populations:** However, adding new features may not improve flowMagic's accuracy for gate boundaries determined by external biological information. Examples include CD161+CD8-, plasma cells, immature neutrophils and CD16-CD14+ populations. These rare populations showed low micro F1 scores (< 60%) even using the template model. Finding training features to improve accuracy for these populations remains an unsolved problem, possibly because they lack recognizable density features, shapes or regular patterns to learn.

**Considerations for more complex models:** While more complex models like Region-based Convolutional Neural Networks (R-CNN) could be considered for future versions of flowMagic, they may not solve the issue of finding an appropriate feature set. These models, while able to generate their own ideal feature maps from pixel input, are more complex to train and show a higher risk of overfitting. Moreover, for inconsistent gates typical of rare populations, which are often determined by external biological information, the automatic estimation of an ideal feature set may be impossible due to the lack of intrinsic data information useful for predicting gate boundaries.

**Potential solutions:** One possible solution could involve changing how the algorithm is executed. Instead of predicting boundaries for complex populations, the algorithm could focus on predicting boundaries for easier populations and indirectly calculating event numbers for more complex ones. For example, it could gate only mature neutrophils (with boundaries directly related to density information) and extract the threshold for immature neutrophils from these boundaries.

Another approach could involve generating different sets of templates for rare populations. For instance, one template could report only mature neutrophils (larger, easier-to-gate populations) while another reports only immature populations (rarer, harder-to-gate populations), focusing the algorithm on generating complex boundaries. However, this strategy would only be applicable if the user has a prior idea of population boundaries.

**Comparison with other approaches.** It's worth noting that while some recent tools like DeepPeak claim high accuracy in identifying rare populations, they often employ multidimensional analysis based on dimensionality reduction techniques. The output of such multidimensional gating tools can be complex for clinical researchers to interpret. In contrast, flowMagic analyzes single bivariate plots, generating output similar to the popular FlowJo tool, which may be more accessible for many users.

### Supplementary Note 5: Data Quality Control (Bot detection)

##### Bots Detection

While in-game rewards increased participation, they also incentivized the use of automated programs (bots) that could automatically submit results, providing higher rewards. These bots generated low-quality results (e.g., a single gate surrounding all data points). The bot detection script (named “user_filter2.R”, see data availability statement link in the main text for Project Discovery bivariate training plots) filtered out results showing gating patterns that deviated significantly from ideal patterns, based on manual review of submitted data (list of review rules is reported below).

1. Submissions that only contain triangles (polygons with 3 vertices).

2. Submissions that only contain two quadrilaterals (polygons with 4 vertices) that have a width larger than 70% of this data range.

3. Submissions that contain out-of-range number of polygons regarding FlowGrid prediction. FlowGrid is an algorithm developed to aid in both detection that estimates the number of polygons based on the kernel density estimation of the x and y axis values of the bivariate plot. For each axis it identifies the number of peaks and possible skewness of the distribution (i.e., asymmetry in the density of the markers values), this can trigger a further partial evaluation to identify possible populations that may be hidden in the 1-dimensional projection. For example, a bivariate that contains two peaks in both kernel densities of the x and y axis and flowGrid will identify if there are 3 or 4 peaks (1 or 2 peaks hidden).

4. Submissions that contain only 1 polygon and the area of the polygon is greater than 90% or less than 10% of the area of the bivariate plot, which is calculated with:

   AreaBiv = Datarangex * Datarangey.

   where:
   - Datarangex= range of data of x axis.
   - Datarangey= range of data of y axis.
   - AreaBiv= area of the whole bivariate plot.

   In other words, this rule removes the situations in which there is one gate around the majority of the events.

5. Submissions that have more than p% of ungated events, where p=13% if more than 2 polygons are present; p=5% if 2 polygons are present; and p=10% if 1 polygon is present. The cutoff percentile follows the general trend observed in the dataset.

6. Submissions that contain polygons that have more than 10 events but less than 2% of the total events number. The 10-event rule is to reconsider players' files that contain good-quality polygons except the ones that have less than 10 events. In this situation, if a polygon contains 5 events, the user file will not be considered a bot solely on this because this polygon will be discarded in the next preprocessing stage.

7. Submissions that only contain quadrilaterals and have more than 10% of the ungated events.

8. Submissions that contain polygons with vertical or horizontal edges on identical x or y axis values. If two vertical edges from different polygons have the same x axis value, the user file is considered a bot because human users should not be able to achieve such accuracy on manual gating and the general trend observed is that players files with this scenario are of bad quality.

9. Submissions that contain a polygon of more than 1000 events with the following features (the peaks are defined with FlowGrid):

   - Polygon width greater than 70% of the x-axis data range and contains more than one peak.

   - Polygon contains two peaks on the x-axis, where each peak's number of events is within the range of 13%-87% of the total number of events within the polygon. The less populated peak has more than 10% or 6% of the total number of events of the bivariate, depending on whether the more populated peak has more than 50% of the total bivariate number of events or not.

   - Polygon contains two peaks on the y-axis, where each peak's number of events is within the range of 12%-88% of the total number of events within the polygon. The less populated peak has more than 10% or 5% of the total number of events of the bivariate, depending on whether the more populated peak has more than 50% of the total bivariate number of events or not.

10. Submissions that contain only 1 polygon and the number of gates predicted by FlowGrid is not 1, or the level of confidence is less than 65%.

The input files of the bot detection algorithm consisted of the data containing the bivariate expression of the events (the data the players were analyzing) and the gating results of each player that gated a specific bivariate plot, each as a single file.

When the situation described in the rule occurred, the researcher established that the result could not be generated by a human player. The researcher also evaluated the quality of the gating process, identifying gates very distant from the ideal situation described before. Before adding the rule to the algorithm, the suspicious plot was visually analyzed by other researchers of the data analysis team to confirm the bot categorization. Hence, the algorithm filtered out the following files submitted through the server system, separating real player files from bots.

**Results of QC Pipeline**: The QC pipeline classified 94% of inputs as bots and 6% as "likely human".

### Supplementary Note 6: Consensus Generation

**Code location.** The script for consensus generation (named “consensus_analysis.py”) is stored with the Project Discovery training plots as described in the data availability statement (https://www.frdr-dfdr.ca/repo/dataset/fe62f923-d7f4-4fd8-9ff1-5bcd9e6f519b). The same link also stores the README file that explains how to run the code.

**Rationale.** While aggregating annotations from multiple annotators is common practice in crowdsourcing[27], some argue that this process may lead to loss of valuable information[28,29,31]. However, for our specific purpose of training a machine learning algorithm for automated gating, we deemed consistent ground truth associated with the same features in the training set necessary[30, 32]. In other words, the curated training set contains only one set of gates associated to each bivariate plot (avoiding multiple set of gates for the same bivariate plots).

Consensus generation addressed three key aspects:

1. Dataset variability: Consensus gates increased training set diversity by avoiding the presence of similar gates generated by multiple players.
2. Consistent truth-pattern association: Machine learning algorithms require consistent input-output pairings for effective training. Inconsistent answers for the same bivariate plots could lead to underfitting due to the model's inability to learn correct associations based on inconsistent information.
3. Mitigation of individual bias: While selecting a random player's answer could achieve the one problem-one answer scenario, it would introduce individual bias. By combining multiple answers, we aimed to generate the best possible answer with broad consensus, avoiding both inconsistency and individual bias in the training set.

**Initial Grouping and Comparison.** Plots that passed bot detection were grouped based on their number of polygons (i.e., the gate boundaries generated by players). Specifically, plots with the same number of polygons for a given bivariate plot were grouped together.

**Selection Criteria and Overlapping Gates.** For further processing (consensus generation), we retained the group of plots meeting these criteria:

1. Highest number of polygons
2. Gated by at least 6 players
3. Mean F1 score of 90% or higher

This process eliminated overlapping gates from the dataset, a crucial step as post-gating biological analysis relies on event counts that must belong to only one gate by convention.

**Consensus Label Generation Algorithm.** The consensus labels algorithm was developed using Python. The code is inside the script file named “user_filter2.R” (https://www.frdr-dfdr.ca/repo/dataset/fe62f923-d7f4-4fd8-9ff1-5bcd9e6f519b). The process involved:

1. Convert player results to a 50x50 pixel map.
2. Assigning weights to pixels according to the number of events within them.
3. Generating consensus using the SpectralClustering algorithm from the scikit-learn library. The SpectralClustering algorithm assigns a label to all events.
4. Discerning labels that represent a proper gate and labels associated with background events. Thus, the algorithm associates all background events with the "0" label (renaming the original label). Events were considered background and associated with the 0 label if they satisfied 2 out of 3 of the following requirements:
   - The original label is associated with a maximum of 3 out of the 4 corner pixels.
   - The original label is associated with the most sparse pixels, meaning the pixels have the greatest distance from each other.
   - The original label is associated with the least number of events.

**Data size**: The resulting consensus dataset occupies 22 GB of storage.

**Labels consistency.** Since each label indicates a different unique gate, it was necessary to assure the labels' consistency across all plots. In addition to training on the density features, the flowMagic algorithm also trains on the coordinates of the events (i.e., markers expression values). Each label is associated with events in a specific position in the plot. An inconsistent association between labels and position would negatively affect the flowMagic algorithm. For example, all events in the same relative position across the plots would be interpreted to be associated with different gates.

On the debotted dataset, the labels were corrected to assure the labels consistency based on the relative location of each polygon (gate) label. The labeling follows these rules:

1. Top left polygon will be labeled first.

2. Left-most position is prioritized more than the top-most position.

The rules can be interpreted visually with a 50x50 grid structure. The bivariate plot is divided into a 50x50 grid and the labeling starts with the top left grid and then moves down before starting on the top of the next column on the right. Within each grid, there are 25 spatial points of equal distance from each other. Throughout the iteration of 2500 grids, the polygon that covers at least 5 spatial points will be labeled first. The iteration ends once all polygons are labeled.

### Supplementary Note 7: Pre-processing, training, prediction and post-processing

**Overview.** The flowMagic algorithm consists of four main steps: input loading, pre-processing, gating (training and prediction), and post-processing. Supplementary Figure 18 provides a visual summary of these steps.

**Input Loading.** flowMagic requires two types of inputs:

1. **Trained model**: A machine learning model trained on flow cytometry (FCM) bivariate data.
2. **Ungated data**: The data to be automatically gated.

The algorithm supports two model types:

- **Template model**: Trained on bivariate FCM data with the same marker combination and hierarchical level as the dataset to be analyzed.
- **Generalized model**: Trained on bivariate FCM data from various marker combinations, panels, and hierarchical levels (e.g., Project Discovery data).

Users should follow this workflow:

1. Extract bivariate data from the first gating step of the hierarchy.
2. Choose between template or generalized model.
3. For template models, optionally use flowSim to select training examples.
4. Apply the chosen model to gate the bivariate plot.
5. Repeat steps 1-4 for each gating step in the hierarchy.

**Pre-processing.** The pre-processing step includes:

1. Quality checking of input files (see pre-processing section below).
2. Data normalization to 0-1 range for each marker (see pre-processing section below).
3. Feature extraction: 34 density features (17 per marker) and 2 expression features, totaling 36 features per CSV file.

**Gating: Training and Prediction.** We evaluated several training models:

- Random Forest (RF)
- Feed-forward Neural Network (FNN)

- K-Nearest Neighbor (KNN)
- Decision Tree (DT)
- Naive Bayes (NB)

The best-performing model was a Random Forest with 2 decision trees, showing 89% cross-validation accuracy and faster processing than KNN. In particular, the training time (considering all repetitions for the tuning parameters) was about 1 minute for the Naive Bayes and Decision tree, 3 minutes for the Random Forest, 4 minutes for the KNN and 32 minutes for the Feed-Forward Neural Network.

Training details:

- Template model: Uses out-of-bag validation.
- Generalized model: Uses repeated k-fold cross-validation (k=2, 100 repetitions).
    - Model A predicts the number of gates.
    - Model B predicts gate boundaries based on Model A's output.

To manage memory constraints (128 GB RAM limit), we implemented a 500-point downsampling for the generalized model during training and prediction.

**Post-processing.** Post-processing aims to resolve overlapping gates, which are particularly problematic in generalized model predictions:

1. Calculate centroids (mean of events) for each predicted class.
2. Compute Euclidean distances between centroids and events.
3. Assign events to the nearest centroid's class.
4. Apply a distance threshold (default 0.15) to prevent fragmented gates.

This process takes 3-4 seconds per plot with downsampling, balancing speed and accuracy.

Supplementary Figure 16 visually summarizes all 4 steps of the flowMagic algorithm: loading of inputs, pre-processing, gating and post-processing.

#### - Loading inputs

The flowMagic algorithm is organized based on the training of template data (i.e, gated data extracted from the single panel and gating hierarchy under analysis) and global data (i.e., gated data extracted from different panels, studies, and gating hierarchies). The FCM data has a hierarchical structure. A gating hierarchy is associated with each biological sample. A single bivariate plot is associated with a gating step (bivariate combination of markers to analyze) of the gating hierarchy. Each bivariate plot can contain multiple cell populations to identify (multiple gate boundaries to generate).

In order to execute flowMagic, the user needs to load 2 inputs:

1) Trained model: ML model trained on FCM bivariate data.

   - The bivariate data is represented as a labeled CSV file. Each row represents the information associated with a single event. The file has three columns: x-axis coordinates (expression of the first marker), y-axis coordinates (expression of the second marker) and classes (reporting the numerical label associated with the event). Each label indicates a different gate of the same bivariate plot (e.g., gate ``1", gate ``2" and so on). The ``0" label indicates background events. Each CSV file is associated with one bivariate plot to be trained. The bivariate data needs to be extracted from a specific gating step of the gating hierarchy. A GatingSet R object can also be provided to automatically generate the labeled data for each training sample of each gating step.

2) Ungated data to analyze: the data to be automatically gated.

   - The bivariate data is represented as an unlabeled CSV file. Each row represents the information associated with a single event. The file has only two columns: x-axis coordinates (expression of the first marker) and y-axis coordinates (expression of the second marker). Each CSV file is associated with one bivariate plot to be gated. The bivariate plot is associated with a gating step of a gating hierarchy. Note that since the data is ungated, the user needs to start from the higher level of the hierarchy. Once the gates of the cell populations of a higher hierarchical level have been automatically identified, the user needs to extract the bivariate data associated with the gating step of the next lower level.

There are two types of possible models depending on the bivariate data trained:

1) Model trained on the template data (template model): ML model trained on bivariate FCM data with the same combination of markers of the same hierarchical level of the dataset to analyze. The bivariate data is extracted from the gating hierarchy of the dataset to analyze.

2) Model trained on the global data (generalized model): ML model trained on the bivariate FCM data containing different combinations of markers, different panels and different hierarchical levels. The bivariate data is extracted from the gating hierarchies of different datasets (Project Discovery data is an example of global data).

The template data focuses the gating on the specific gating patterns that the user considers significant. The FCM dataset to be gated may be extremely heterogeneous. The data to gate may be different from the bivariate example trained by the user. To compensate for this disadvantage, the users need to provide multiple bivariate examples in the template data. In other words, the template data may be associated with the bivariate data of a single bivariate plot (single labeled CSV file) or to the bivariate data of multiple bivariate plots (multiple labeled CSV files). Determining how many examples to consider for the training of the template data is fundamental for the accuracy of the gating. It is also complex to evaluate because it depends on the cell population to analyze. The application of the flowSim tool represents a possible approach to select the ideal set of examples to be included in the template data. The flowSim tool is designed to identify the most heterogeneous examples in a bivariate FCM dataset. Thus, the user can train on the bivariate examples that represent the most heterogeneous component of the dataset to analyze (see Discussion).

In general, based on which model is used as input, the user needs to follow this workflow:

1) Extraction of the bivariate data to be gated from the first gating step (the first bivariate combination of markers represented in a bivariate plot) of the gating hierarchy (highest level of the hierarchy).

2) Choose input model to use: template model or generalized model.
    - If the user wants to apply a template model, executing flowSim on the extracted bivariate data can suggest what examples to include in the relative template data. Once the training examples have been identified, the user uses the flowMagic training function to perform the training of the selected examples, generating a template model. The user can choose for training a subset or all of the examples selected by flowSim. The user applies the trained model to gate the bivariate plot of this gating step. The user can also ignore the flowSim step choosing a random example to be used as a template.

    - If the user wants to apply a generalized model, the flowMagic package already provides a generalized model trained on the PD dataset. It is sufficient to load this model to gate the bivariate plot of interest

3) Step 1 and step 2 are repeated until the user has completed all the gating steps of the hierarchy. At each repeated step 1, the user extracts the data associated with the combination of markers and events of the next population to gate (in a lower hierarchy level) according to the gating hierarchy.

#### - Pre-processing

The input CSV files are loaded after a quality check offile format, as reported by the flowMagic package documentation (e.g., checking that the files contain the two columns of marker expression). For each marker, the data is normalized to a 0-1 range and the following features are extracted: number of peaks (peak=local maximum), height of each peak, position of each peak, and start and end of density changes surrounding each peak. Considering a max of 4 peaks per marker, there are 17 density features for each marker, 34 density features in total. The expression of each marker is also used as a feature, generating a total of 36 features during the pre-processing of each CSV file. The final training dataset is composed by the extracted features of all input bivariate examples (all input CSV files). The same set of features is generated for the data to gate.

##### - Gating step: training and prediction

In our performance evaluations tests, the A078 dataset was used as a training dataset to generate the generalized model. Both the final A078 dataset with consensus gates (31,703 files) and the A078 without consensus gates (575,396 files) were trained and tested. The template model was generated based on the bivariate data extracted from each dataset tested. The training, cross-validation and prediction/testing step were performed using the caret R package. This is the most popular package to perform machine learning using the R language, providing a framework that optimizes the computational time and memory load.

Several different training models were tested using the same training features described before: Random Forest (RF), feed forward neural network (FNN), K-nearest neighbor (KNN), decision tree (DT), and Naive Bayes (NB). For each model, 5 different values for each parameter were tested. The number of trees (n.trees) in the random forest, the decay and the number of units in the hidden layer (size) for the neural network, the probability distribution of the classes (kernel estimate or gaussian distribution) in the Naive Bayes, the k clusters in the KNN, and the complexity parameter (cp) in the decision tree. The details for each parameter can be found on the caret R package documentation. All models were tested using a repeated k-fold cross validation method with k=2. The events of 50 random PD bivariate plots were used as a training set and the events of 1 random PD plot as a validation set for 10 repetitions. The tests were performed on a dataset composed of 500 random PD plots. The best training model was a random forest composed of 2 decision trees, showing the highest cross-validation accuracy (defined as the mean percentage of correct predictions across all validations sets and repetitions) and training speed (about 20 seconds). Supplementary Table 1 reports the best cross-validation accuracy for all models and their best parameters.

Once the best model to train was established, the template model and the generalized model were trained. Note that the template model needs to be trained and validated for each gating step to analyze, while the generalized model needs to be trained and validated only one time and applied directly on each gating step. The out-of-bag validation method is the default validation method for the training of the template model.

In the case of the generalized model, two types of models are trained. One model is trained to predict the number of gates (“Model A"), with 6 gates being the maximum number. The second model is trained to predict the gates’ boundaries (“Model B") based on the predicted number of gates. There is a Model B for each possible number of gates present in the training set. For example, Model B.2 draws two polygons (gates boundaries associated with two gates), Model

B.3 draws 3 polygons and so on. During the automated gating, after Model A predicts the number of gates, the associated Model B draws gate boundaries. Note that the distinction between the two models (Model A and Model B) is not necessary in the template model previously discussed as the number of gates is pre-defined by the templates (only the gates' boundaries are needed in these cases). A repeated k-fold cross validation method with k=2 is the default validation method for both GP models. The number of examples used in each k part depends on the two types of models generated. For Model A, the events of 5,000 random PD examples were used as a training set and 10 random PD examples as a validation set for 100 repetitions. Each Model B is trained on the subset of examples having the same number of gates (e.g., Model B.2 is trained only on plots having 2 gates). A maximum of 3,000 PD random examples were used as a training set and 1 random PD example as a validation set for 100 repetitions.

By default, when training the generalized model, a downsampling of plots is performed (500-point downsampling is the default) to limit memory load and address possible class imbalance (i.e., different number of points among the gates). Note that the testing was limited by the memory constraints of the machine used to perform the analysis (128 GB RAM, Intel Core i7 Processor). When training using 500-point downsampling, the memory load is below 10GB. With no downsampling, the memory load is above the machine limit of 128 GB. Note that the machine used for testing has a memory setup superior to the average desktop computer (which is about 8-16 GB in the last decade). More training iterations and more training plots than the ones used in the testing described here may improve the generalized model accuracy, but this testing will require a memory superior to 128 GB. A possible solution to this memory limit may be the usage of a web cloud service, like Amazon Web Server (AWS) cloud service or similar services, with a virtual machine having a memory of 500 GB or above.

The downsampling is also performed during the prediction step (of the generalized model) to be consistent with the downsampling performed during its training step. In addition, the generalized model predictions always require a complex post-processing step that slows down the processing of the input data; thus, the downsampling speeds up the prediction step execution (see post-processing section).

By default, there is no downsampling for the template model since it is faster, less memory demanding and more accurate than the generalized model in all situations including class imbalance situations. Of note, in case of class imbalance situations for the template model, the user can just generate a template designed only for the smaller cell population (only 1 class to gate), nullifying the imbalance among the classes. Of course, this strategy cannot be used for the generalized model that trains on all classes in all bivariate plots at once. In order to speed up execution (e..g, when gating bivariate plots having a high number of events), it may be possible to downsample for the template model. Because of the need to reduce the memory load, the training of the dataset without consensus plots was performed by training a different random subset of plots in two simulations. Each subset is generated keeping the same number of plots for each number of gates (for example: 10,000 plots with 2 gates, 10,000 plots with 3 gates and so on). Plots with a high number gates (like 5 or 6 gates) are under-represented even in the whole original dataset. The generation of a subset contributes to reducing the imbalance. Each subset per simulation contains about 43,000 plots, a larger dataset than the dataset with consensus gates. Each Model B is trained in the same way as the consensus dataset, described previously. In order to reduce memory load, Model A was trained using 1,000 random PD examples for 50 iterations.

#### - Post-processing

ML predictions generated by the generalized model tend to produce overlapping gates. As previously explained, overlapping gates cannot be tolerated as final gates because they alter the post-gating analysis. Overlapping gates are generated when the trained ML model makes predictions that determine concave boundaries with jagged borders. The PD dataset does not contain overlapping gates. In order to avoid overlapping gates, the boundaries need to be smooth and with convex angles. The players were instructed to draw the gates in this way. Cases distant from ideal situations were removed during the data quality checking process. In order to avoid overlapping gates, the algorithm needs to ensure a smooth separation between the events associated with one predicted label and the events associated with the other predicted label. When the variability of the training set is high, the training process is more difficult, and the final predictions may not always generate a smooth separation of the different boundaries, generating jagged borders (Supplementary Figure 17-18).

The ML predictions generated by the template model do not show this effect or it is very limited because the set of examples is less variable than the set of examples contained in the PD dataset. No complex post-processing is necessary for the template model. In order to remove the large overlapping boundaries generated by the generalized model, the centroid of each predicted class is calculated. Then, the Euclidean distance between each centroid and each event is calculated. The events are assigned to the class with the nearest centroid. Based on the number of events and polygons, this process may take several minutes per plot. Thus, a downsampling of the events during the prediction step is necessary. The prediction step downsampling reduces the post-processing time to 3-4 few seconds per plot, but it may also negatively affect the accuracy. The post-processing required the tuning of a parameter that controls the distance at which the events belonging to two different centroids are combined to avoid the generation of fragmented gates. In other words, it regulates the minimum acceptable distance between two centroids (current default is 0.15).

### Supplementary Note 8: ethical data considerations

This study follows the most recent Human-centric computer vision (HCCV) data curation practices to assure the generation of a high quality unbiased dataset. Non-consensual data collection may lead to the generation of biased training datasets difficult to evaluate and with the potential risk of retraction. All the annotators involved in our study (the players of the massively multiplayer online game *EVE Online*) were informed about the future usage of their work before the start of the annotation process. The project was freely accessible and included an introduction page that described the project. The purpose of the project and potential data use were clearly described in publicly accessible web spaces promoted through web social channels. Since the annotation task was associated with a specialized professional sector (cytometry scientific research), in order to assure both the fairness and accuracy of the annotation task, we provided all players with a complete set of instructions and ideal examples that clearly describe the annotation task (i.e., the gating task). The game's complexity attracts a user base composed of adult individuals interested in facing complex challenges. The user base is composed of players from all over the world and with different sociodemographic factors. The annotation task consists of drawing boundaries around a group of points, a natural skill we assumed present in any healthy human being. Therefore, we considered the annotation task itself free from any possible bias potentially caused by sociodemographic factors (such as sex, gender, ethnicity, religion, income or education) or geographical factors (such as country, city or region). The data was collected respecting the anonymity of the users. Only the gates generated by the players and their associated metadata (like age) were collected together to perform the evaluation analysis. Overall, we acknowledged the HCCV practices of fairness, privacy, and consensual collection, as reported in Andrews, Jerone et al. "Ethical Considerations for Responsible Data Curation." 10.48550/arXiv.2302.03629. 2023

### Supplementary Tables and Figures

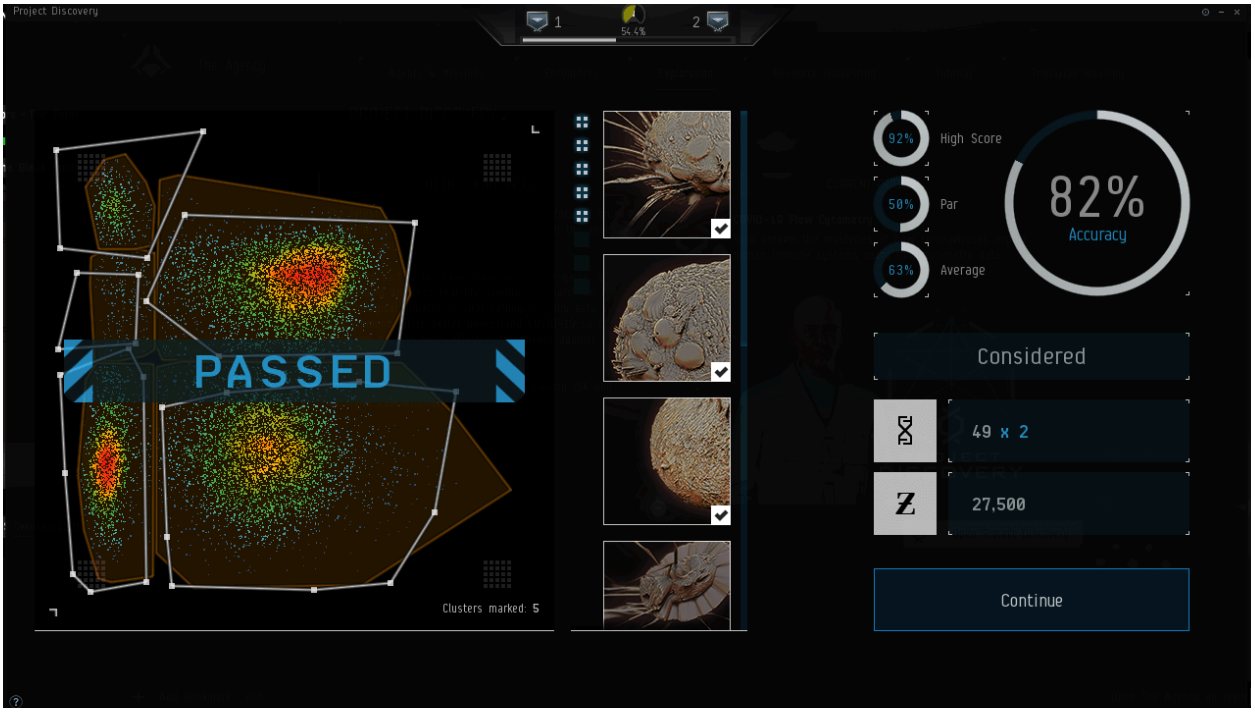


Supplementary Figure 1: **Gating mini-game in *EVE online*. Example of gating performed within the *EVE online* server.** On the left, there is the plot to be gated. The white boundaries represent the gates generated by the players. The yellow areas represent the gold standards for that plot. On the top right, the accuracy value of the gating performed by the users is reported. The in-game rewards are reported below the accuracy value. Plot-specific statistics are also reported next to the accuracy score (e.g., average score and highest score for that plot).

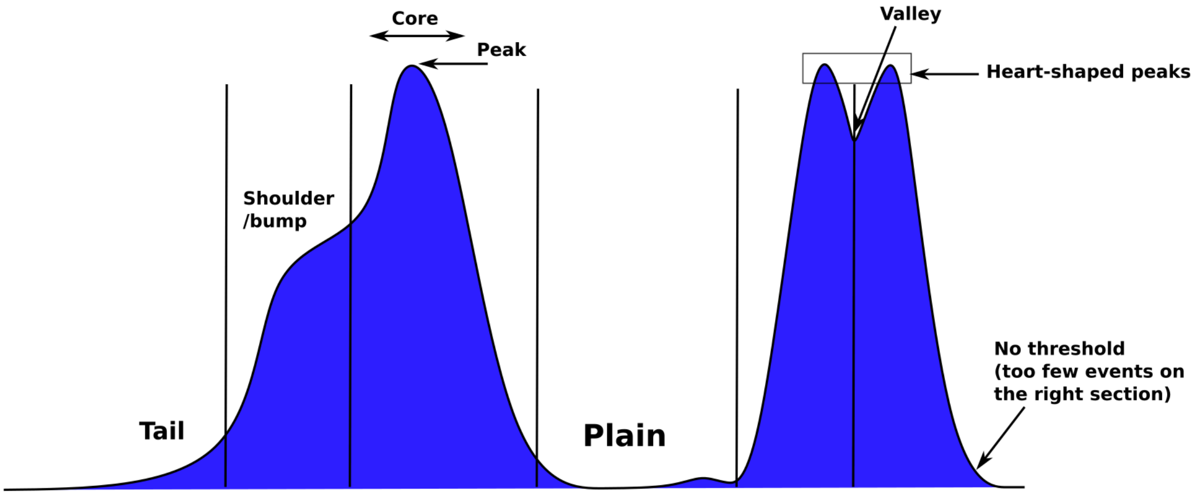


Supplementary Figure 2: **Density elements to determine gating rules for ideal gates.** The density rules are separated by black vertical lines across the density plot. These rules were used to evaluate the quality of the players' gates.

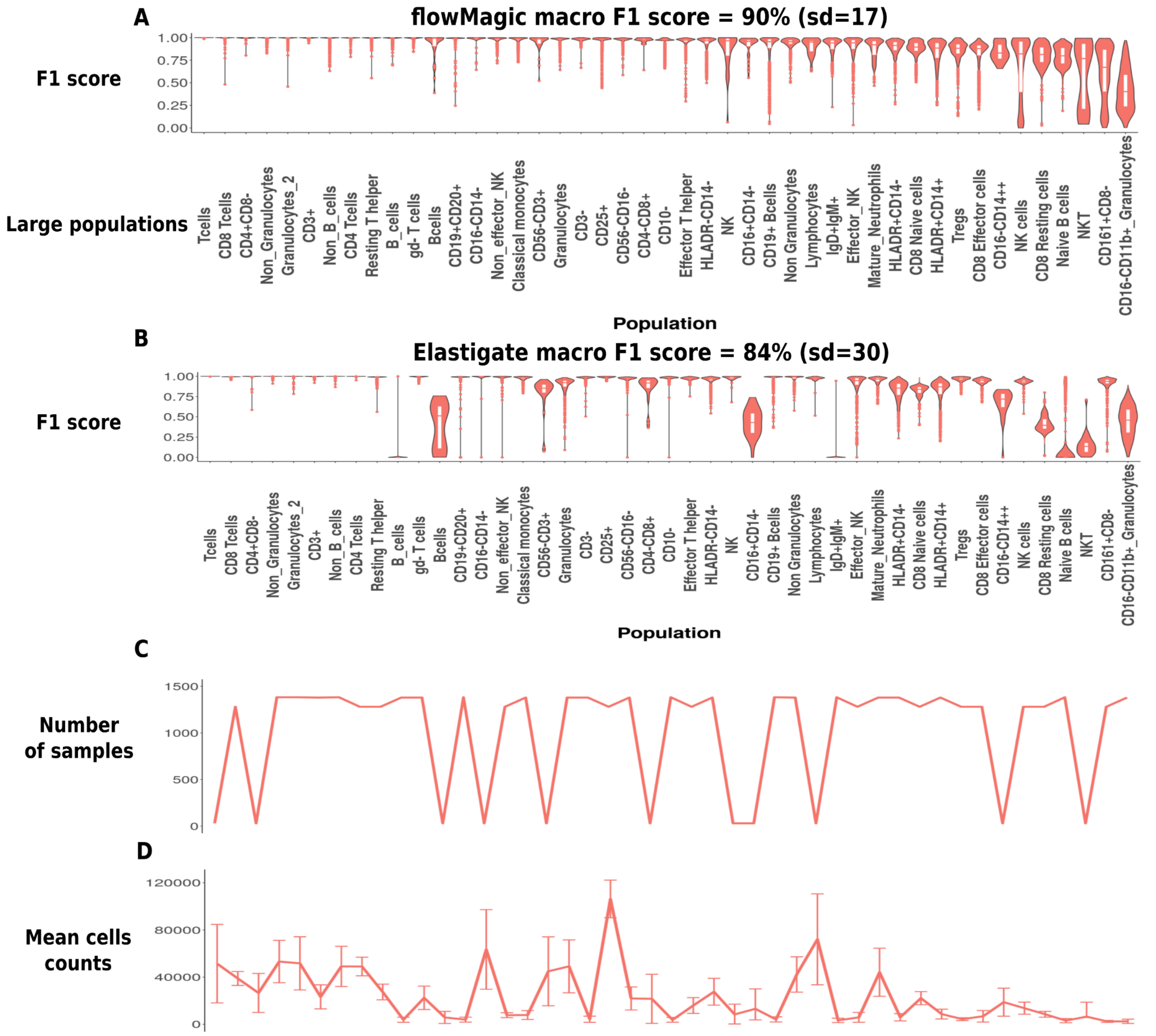


Supplementary Figure 3: **flowMagic and Elastigate results considering one template training for gates having large populations.** (A) flowMagic F1 scores, (B) Elastigate F1 scores, (C) number of biological samples and (D) mean counts for each gate under analysis. ”sd” indicates the standard deviation.

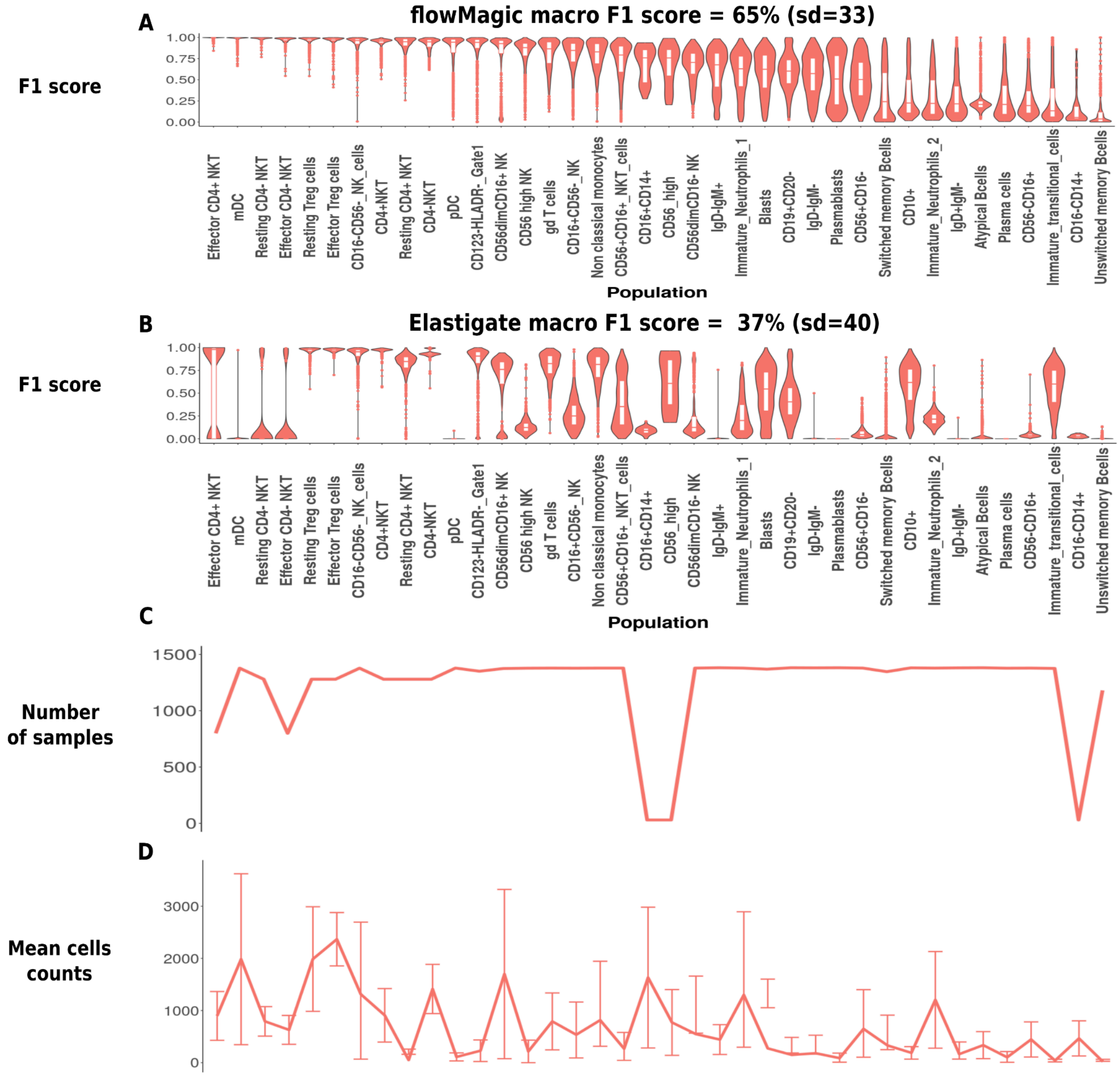


Supplementary Figure 4: **flowMagic and Elastigate results considering one template training for gates having rare populations.** (A) flowMagic F1 scores, (B) Elastigate F1 scores, (C) number of biological samples and (D) mean counts for each gate under analysis. ”sd” indicates the standard deviation.

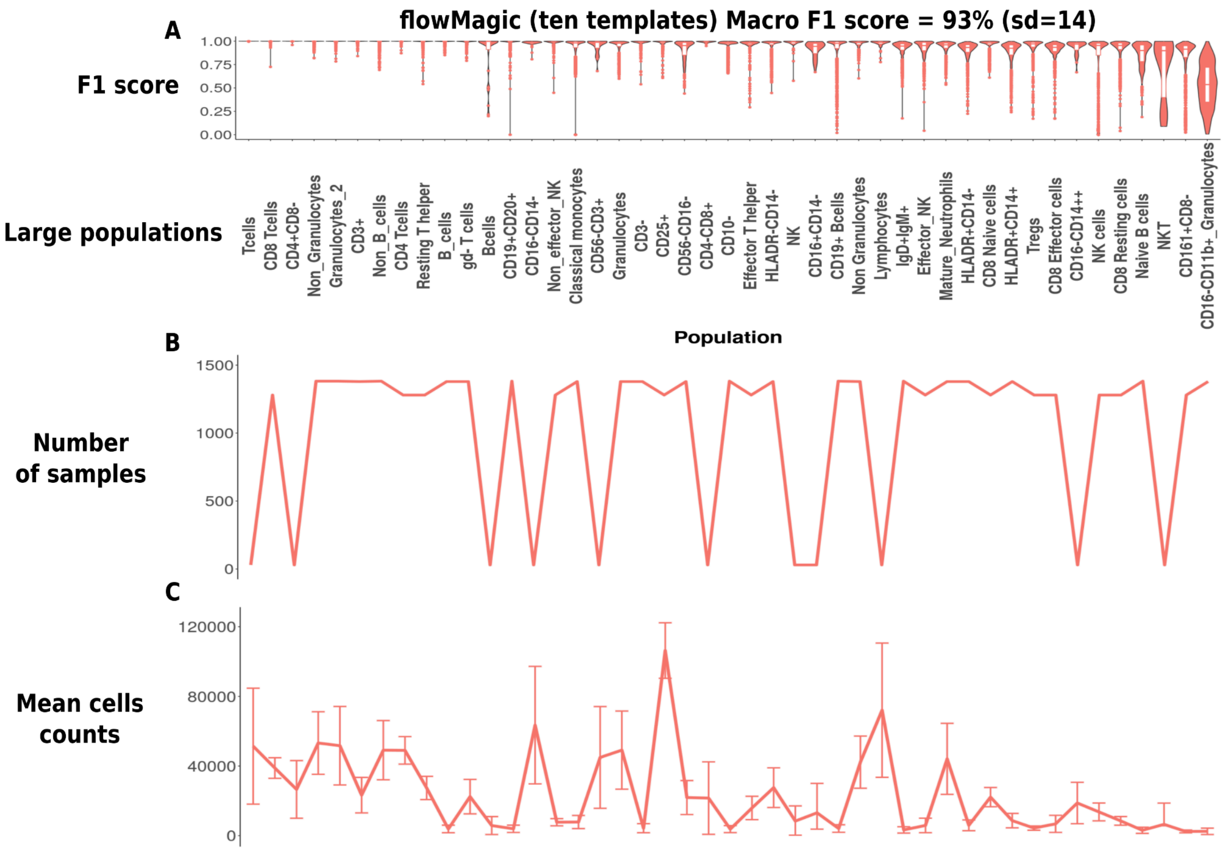


Supplementary Figure 5: **flowMagic algorithm results considering ten templates training for gates having large populations.** (A) F1 scores, (B) number of biological samples and (C) mean counts for each gate under analysis. ”sd” indicates the standard deviation.

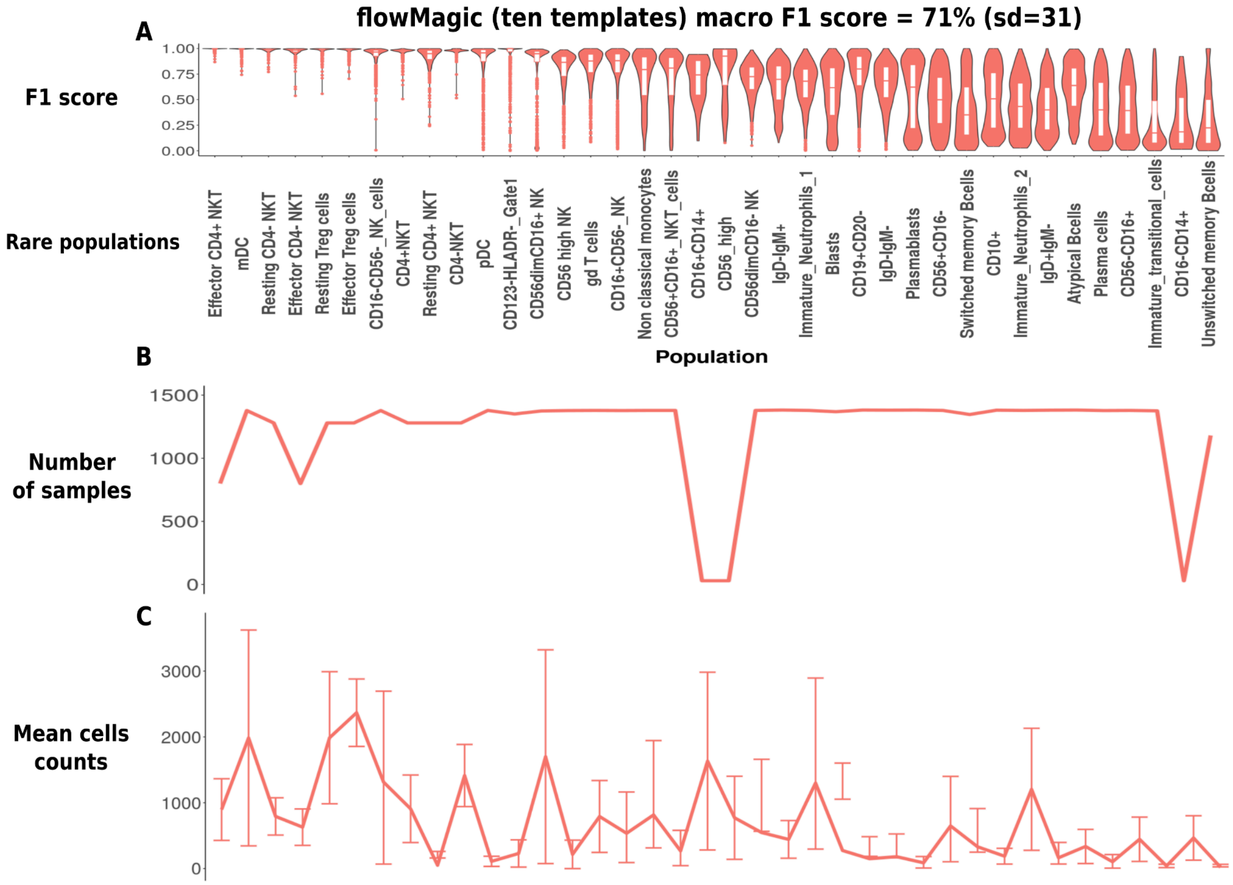


Supplementary Figure 6: **flowMagic algorithm results considering ten templates training for gates having rare populations.** (A) F1 scores, (B) number of biological samples and (C) mean counts for each gate under analysis. ”sd” indicates the standard deviation.

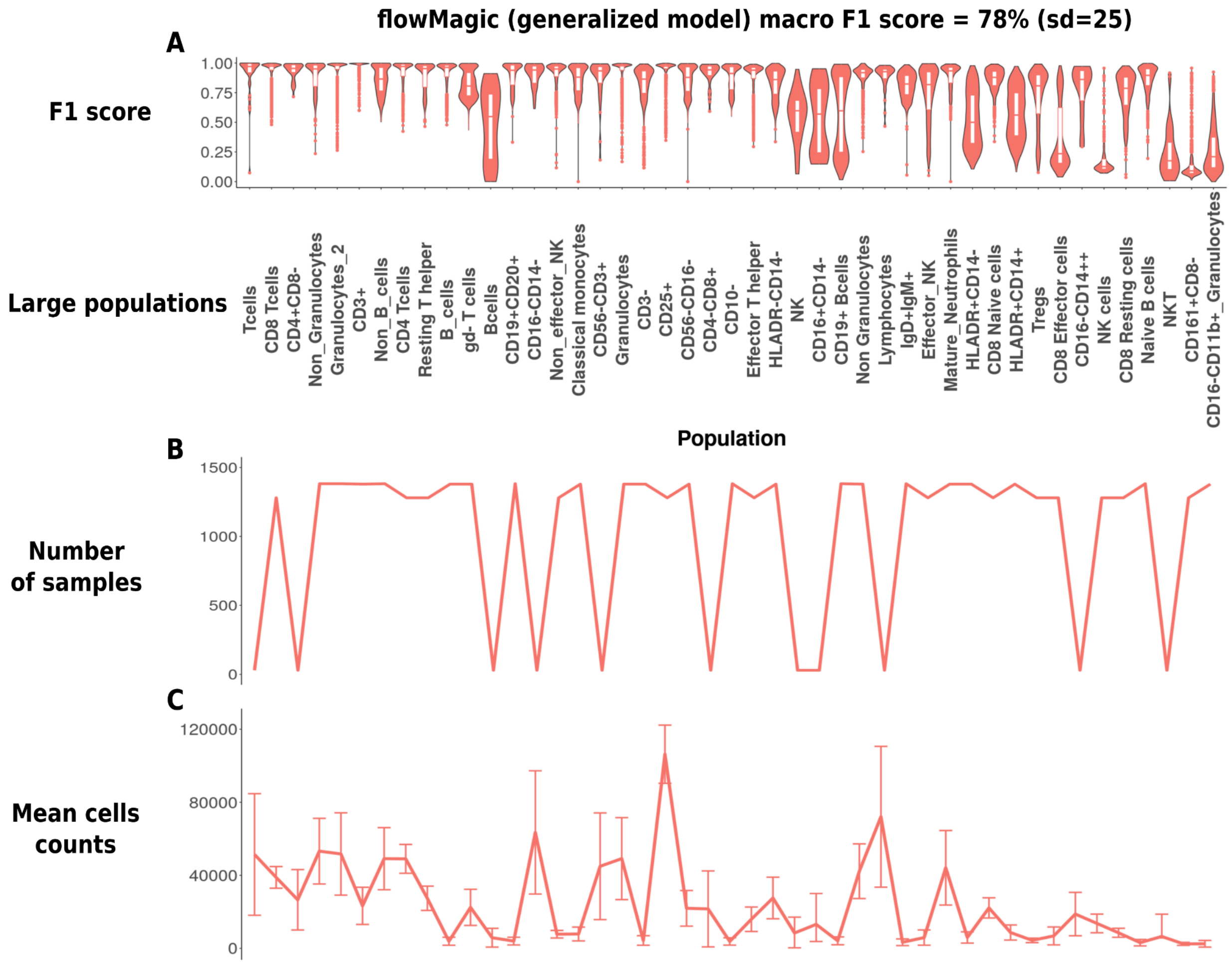


Supplementary Figure 7: **flowMagic algorithm results generalized model for gates having large populations.** (A) F1 scores, (B) number of biological samples and (C) mean counts for each gate under analysis. ”sd” indicates the standard deviation.

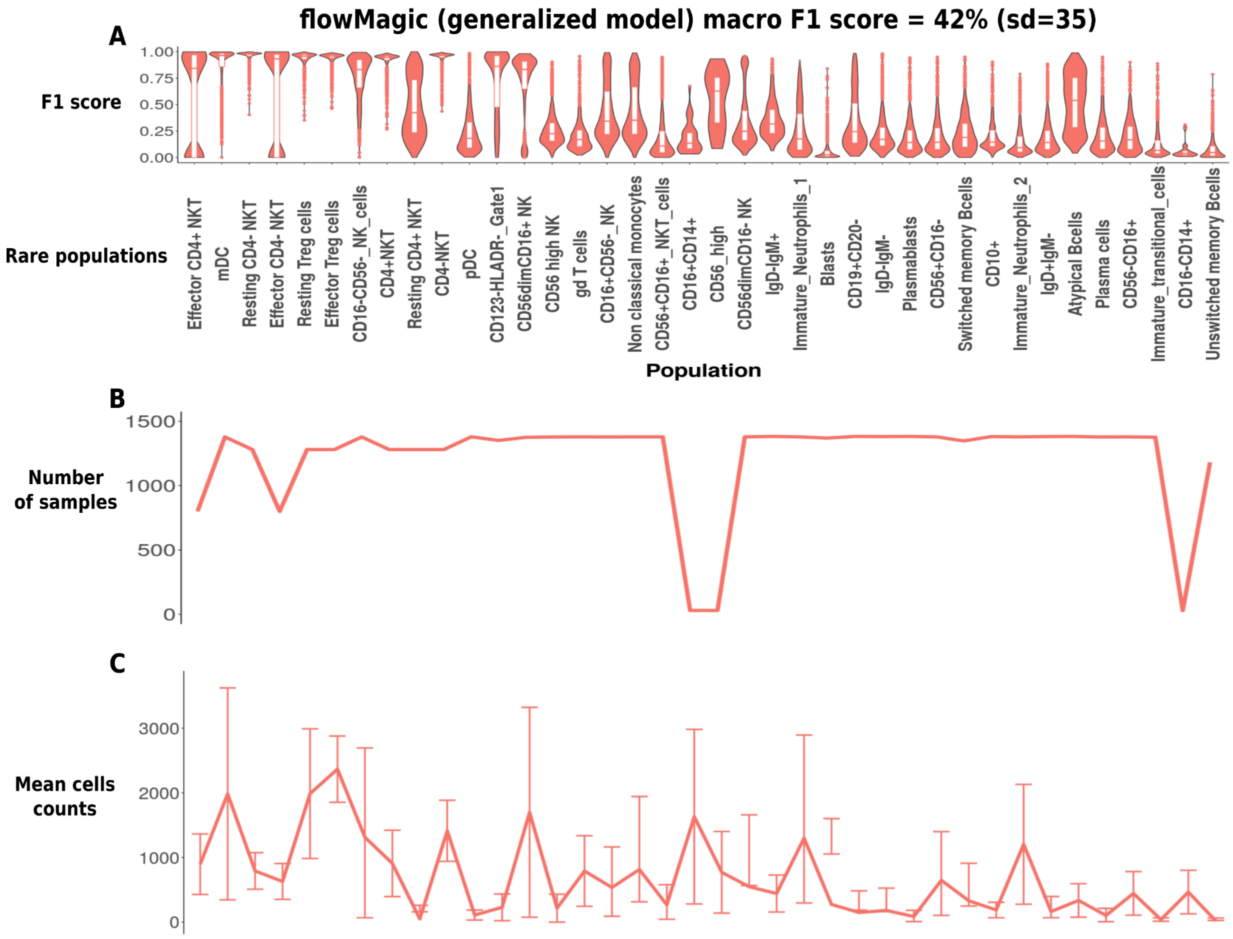


Supplementary Figure 8: **flowMagic algorithm results generalized model for gates having rare populations.** (A) F1 scores, (B) number of biological samples and (C) mean counts for each gate under analysis. ”sd” indicates the standard deviation.

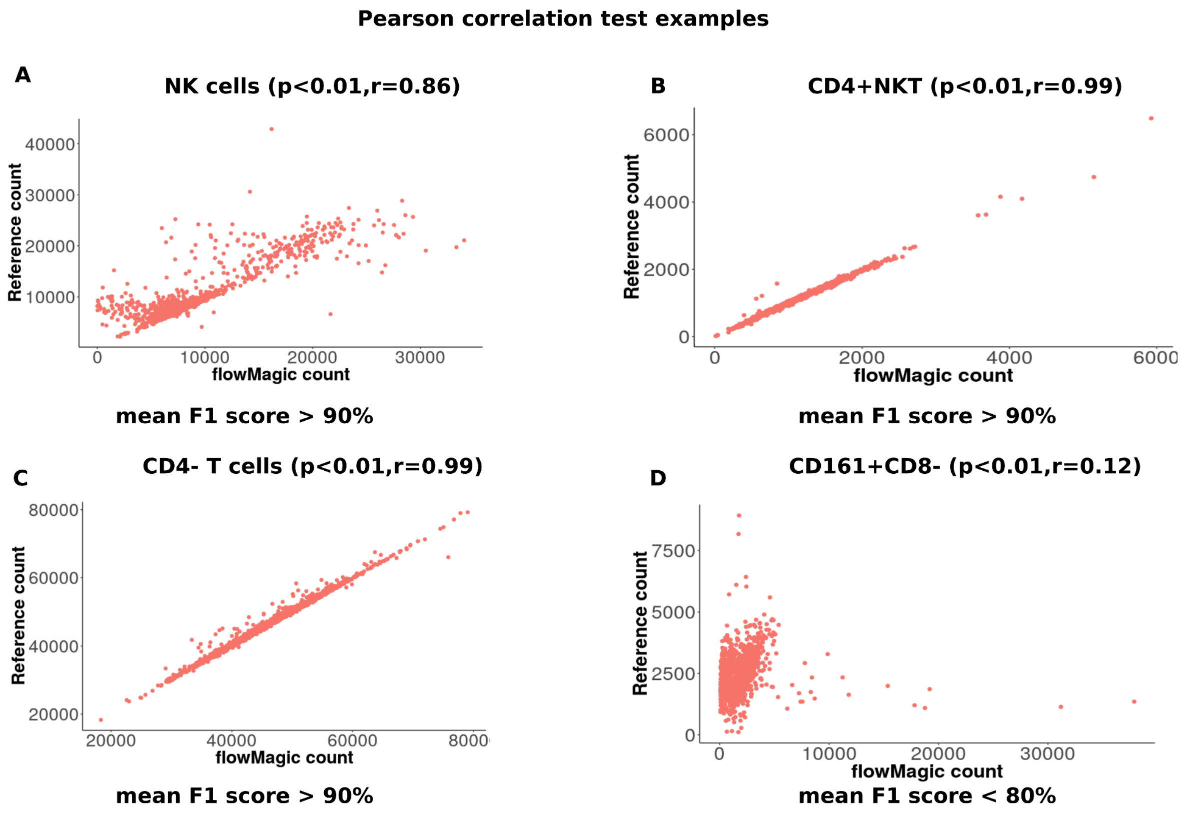


Supplementary Figure 9: **Correlation between flowMagic counts and reference counts using templates of the same centre.** (A-D) Above each correlation plot, the p-value (p) and correlation coefficient (r) of the Pearson correlation test is reported.

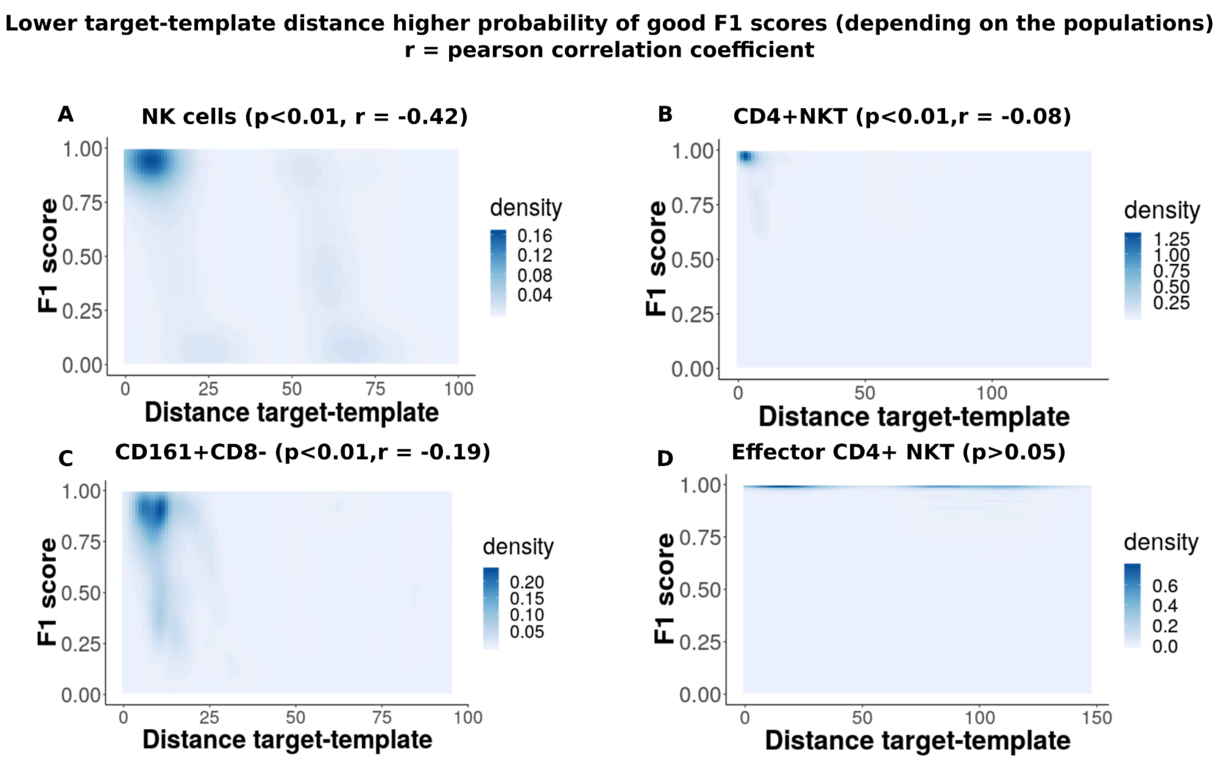


Supplementary Figure 10: **Lower target-template distance, higher F1 scores for most populations.** (A-D) Density scatter plot between F1 scores (y axis) and template-target distance (x-axis) for different populations. Significant correlations calculated using the Pearson correlation test (p<0.05), r=Pearson correlation coefficient.

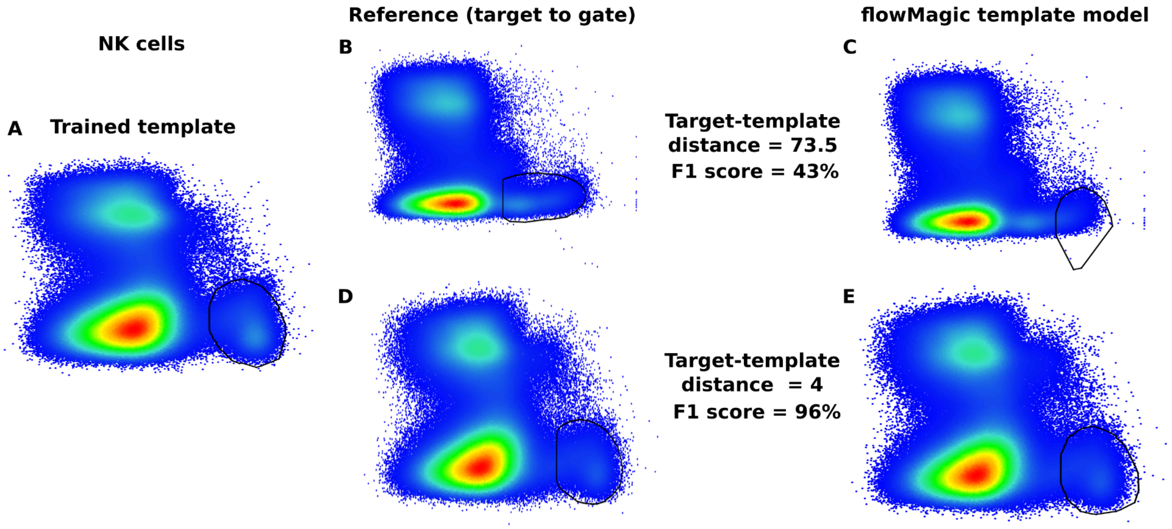

Supplementary Figure 11: **F1 scores of cell populations having inconsistent gates boundaries are negatively correlated with target-template distance.** (A) Trained template used to calculate distance. Reference gates representing the target to analyze and automated gates for (B, C, D, E) two examples with different F1 scores and target-template distance.

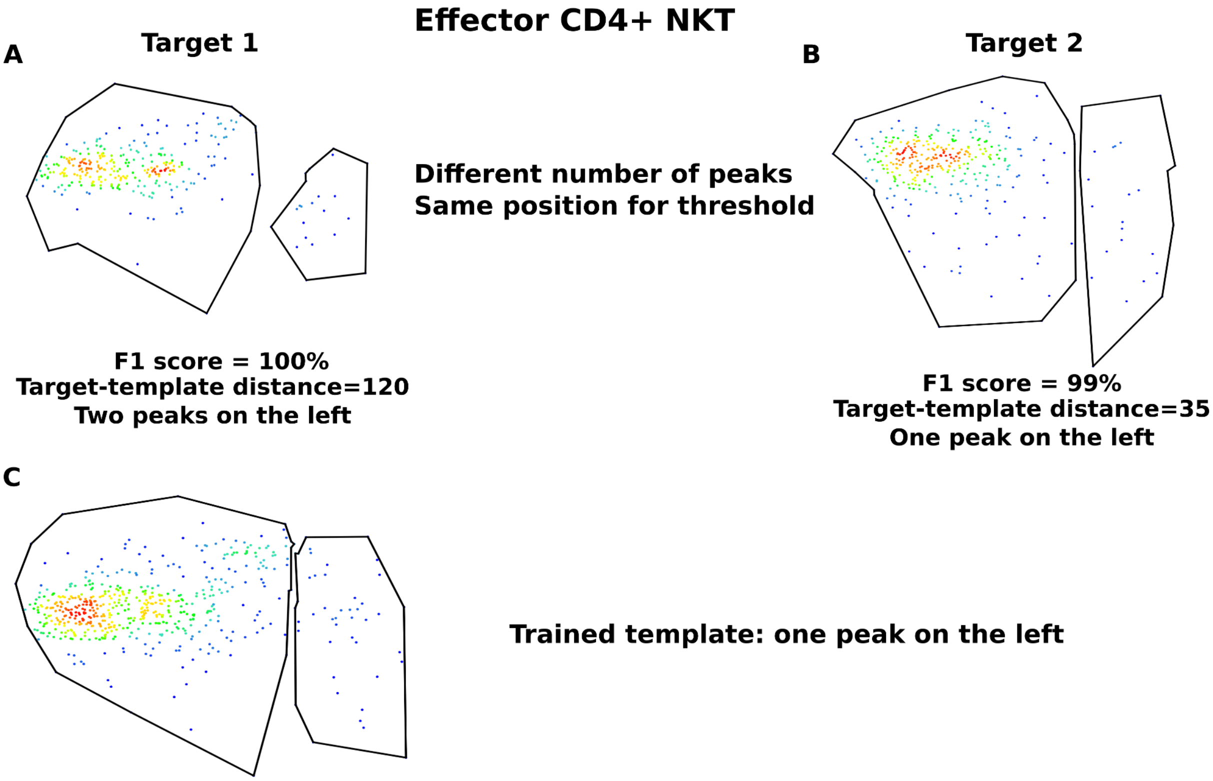


Supplementary Figure 12: **Target-template distance is not correlated with accuracy for populations having consistent gates boundaries.** (A, B) Reference gates representing the target to analyze and automated gates for two examples with different F1 scores and target-template distance. (C) Trained template used to calculate the distance.

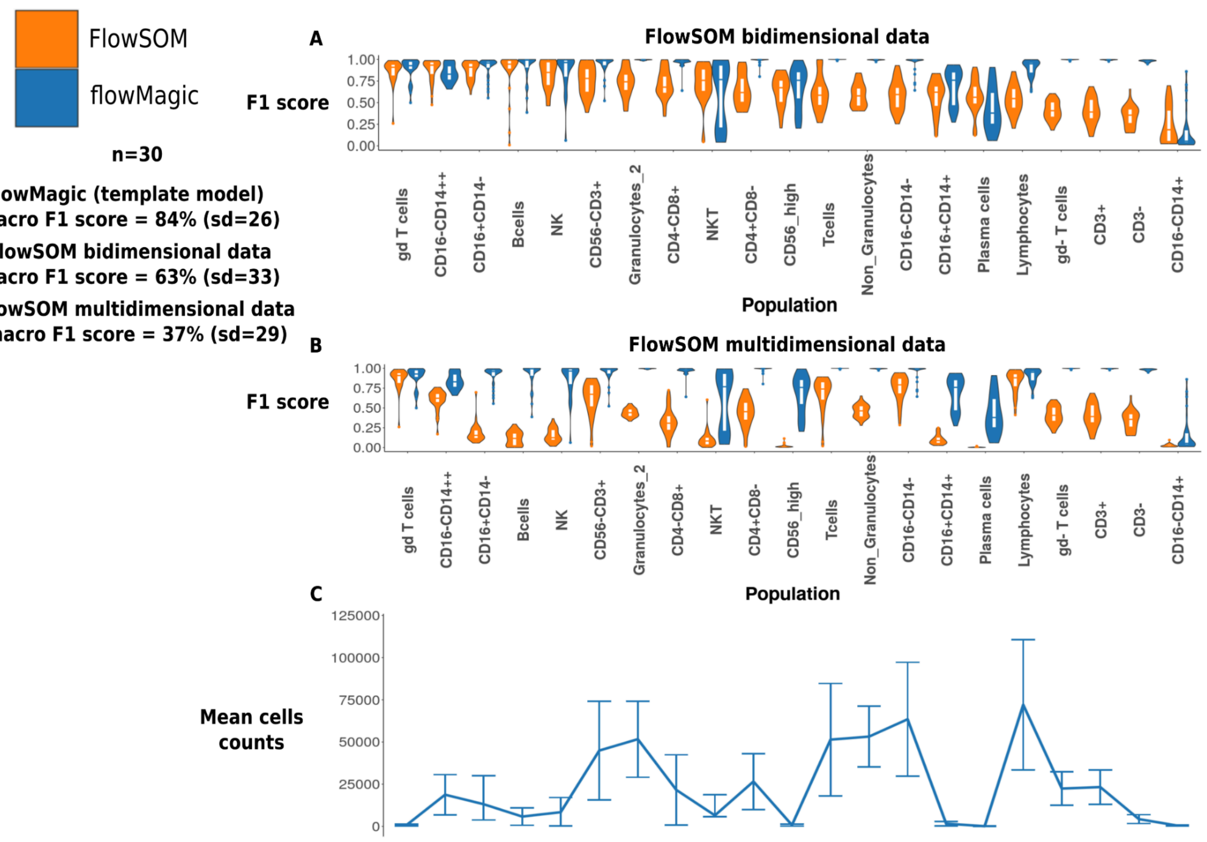


Supplementary Figure 13: **flowMagic (one template) shows superior performance compared to the FlowSOM tool.** (A) F1 scores generated using flowMagic template model (one template) using FlowSOM on bidimensional and (B) multidimensional data. (C) The mean number of counts for the reference gates is indicated below each reference gate label. “n” indicates the number of biological samples tested. “sd” indicates the standard deviation.

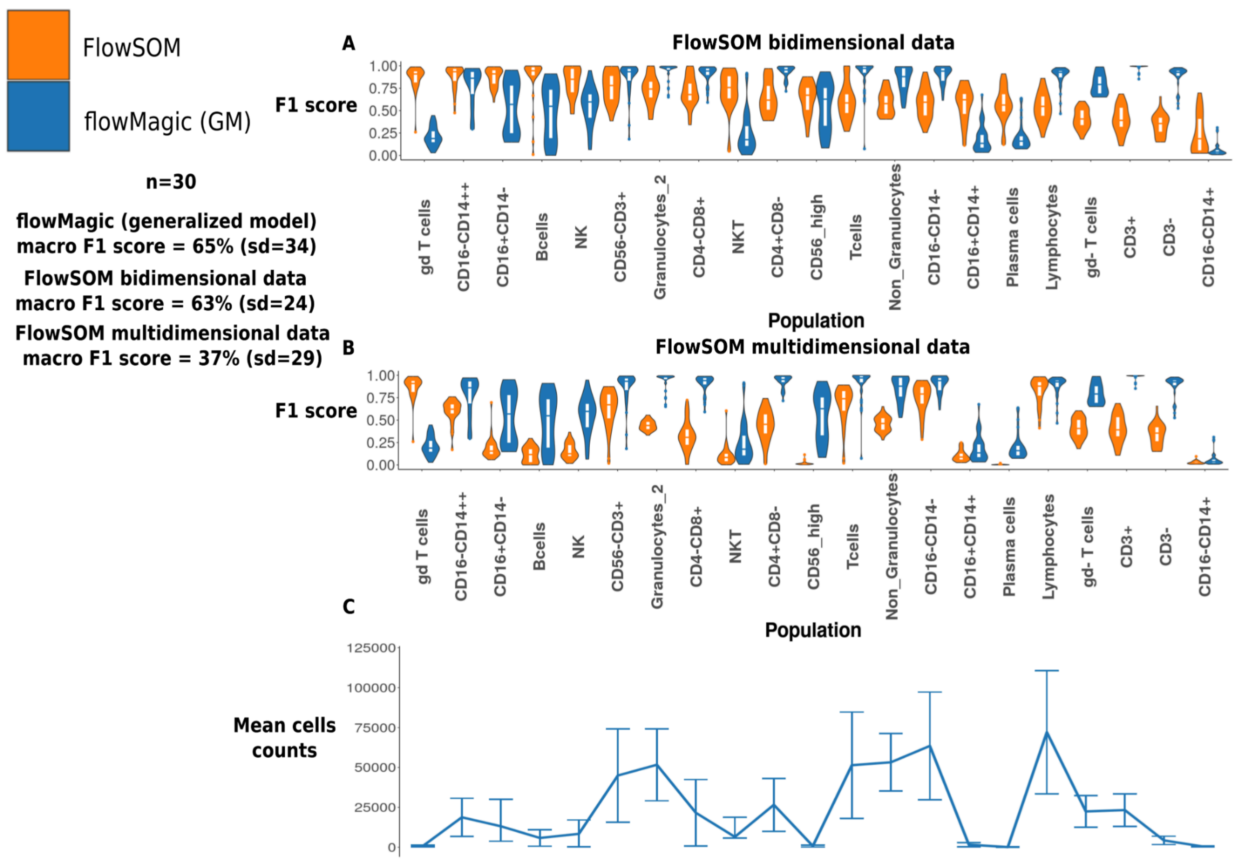


Supplementary Figure 14: **flowMagic (generalized model) shows superior performance than the FlowSOM tool.** (A) F1 scores generated using flowMagic generalized model using FlowSOM on bidimensional and (B) multidimensional data. (C) The mean number of counts for the reference gates is indicated below each reference gate label. “n” indicates the number of biological samples tested. “sd” indicates the standard deviation.

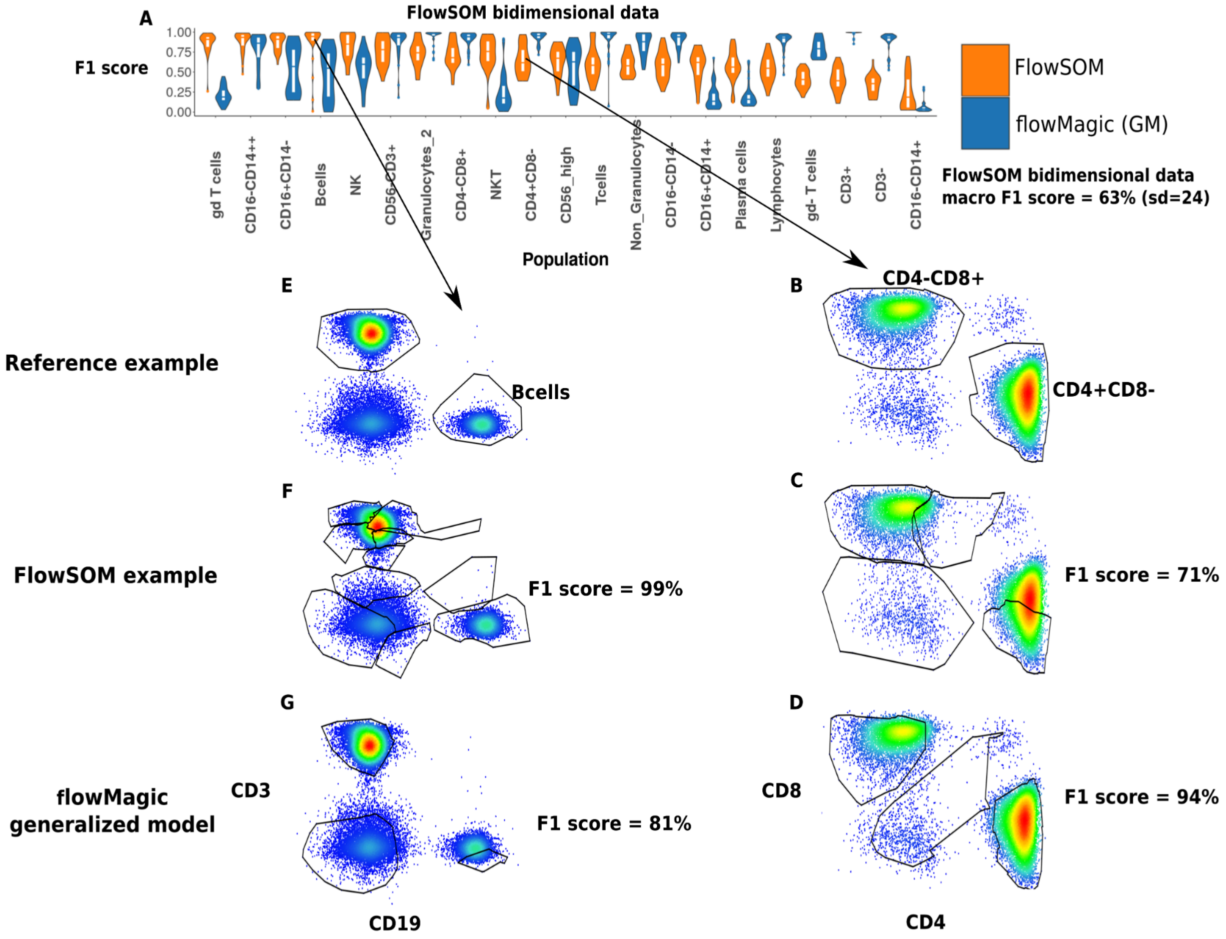


Supplementary Figure 15: **flowMagic (generalized model) and FlowSOM examples.** (A) F1 scores generated by FlowSOM on bidimensional data. (E,F,G) flowMagic generalized model with an example of FlowSOM gate having a higher F1 score than flowMagic and (B,C,D) an example of FlowSOM gate having lower F1 score than flowMagic.

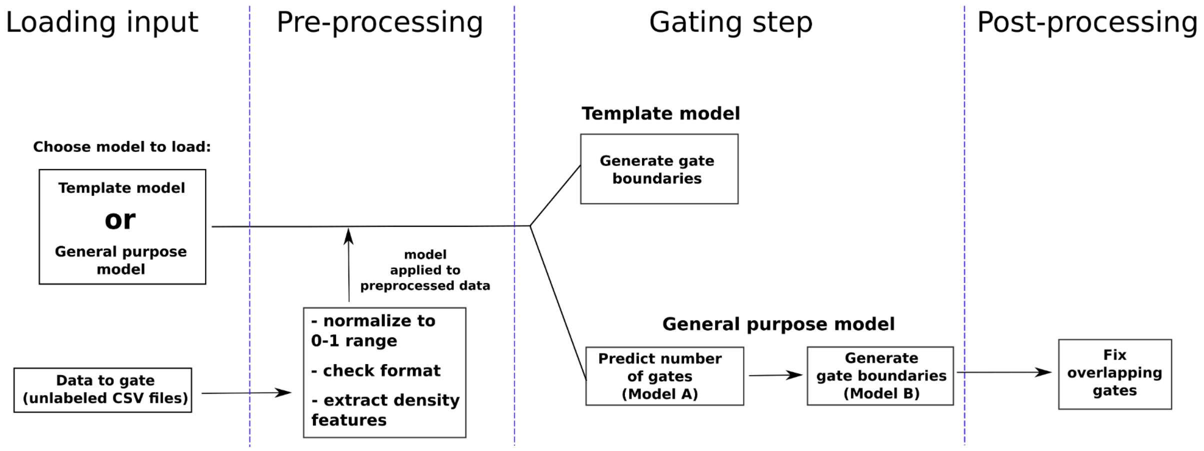


Supplementary Figure 16: **flowMagic flowchart.** The four steps composing the flowMagic algorithm: inputs loading, data preprocessing, gating step, post processing step.

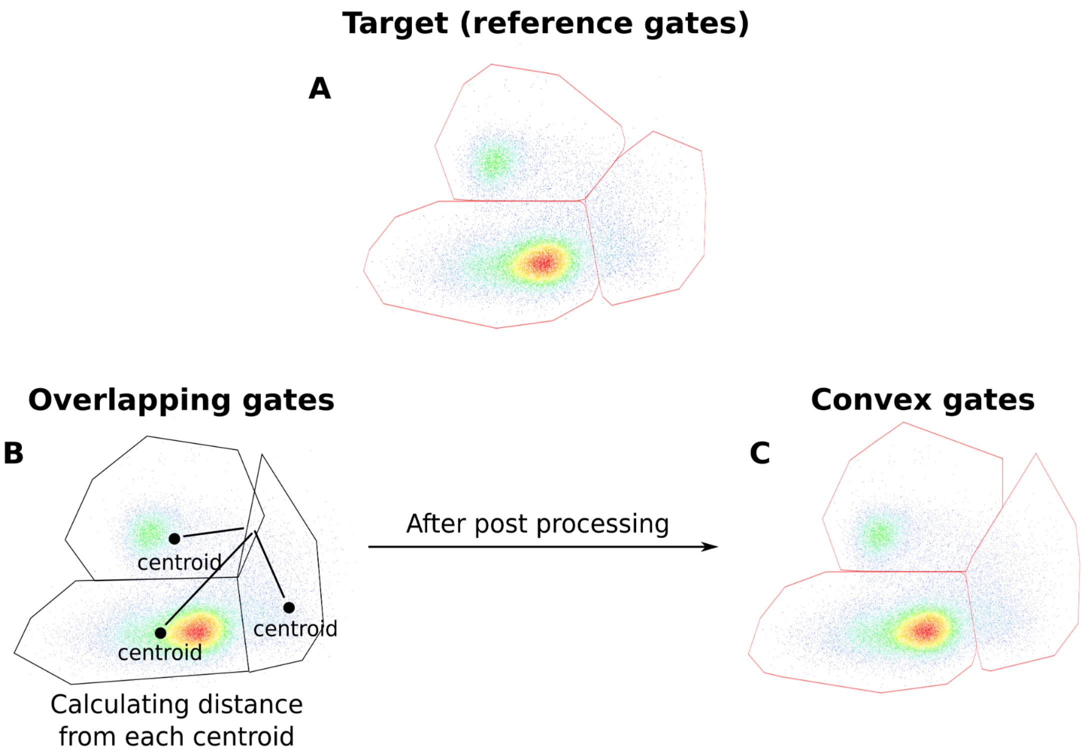


Supplementary Figure 17: **flowMagic post processing to fix overlapping gates.** (A) reference gate; (B) detection and removal of overlapping gates; (C) final convex gates.

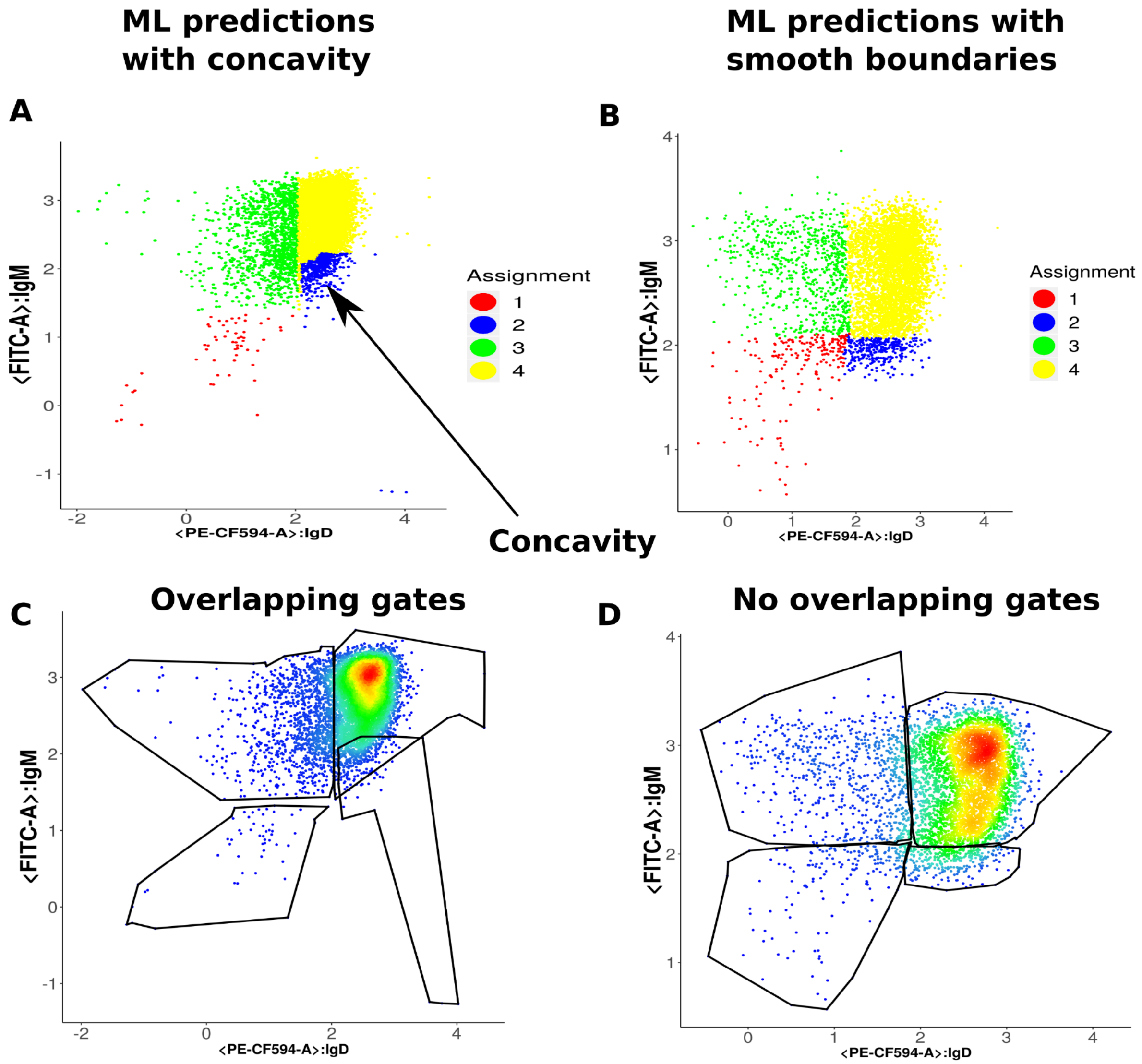


Supplementary Figure 18: **Example of events assignments generating concavity.** (A) Example of concavity with events coloured based on their gate assignment. (B) Example of smooth boundaries with events coloured based on their gate assignment. (C) Example of polygons with overlapping boundaries based on the events assignment shown in (A). (D) Example of polygons with no overlapping boundaries based on the events assignment shown in (B).

Supplementary Table 1: **Both FlowMagic models show superior macro F1 scores than flowSOM.** Comparison of mean macro F1 scores across all populations between FlowSOM (first column), flowMagic template model (second column) and flowMagic generalized model (third column).

| FlowSOM | flowMagic template model | flowMagic GM |
|---|---|---|
| 63% | 84% | 65% |

Supplementary Table 2: **FlowMagic template model shows superior macro F1 scores than Elastigate.** Comparison of mean macro F1 scores across large and rare populations between Elastigate (first column), flowMagic template model (second column) and flowMagic generalized model (third column).

| Size population | Elastigate | flowMagic template model | flowMagic GM |
|---|---|---|---|
| Large (> 2,367 events) | 84% | 90% | 78% |
| Rare (< 2,367 events) | 37% | 65% | 42% |

Supplementary Table 3: **Studies sources.** The first column reports the identifier of the dataset. The second column reports the source of the dataset. The source column reports the link for the original publication that reports the instructions to access the data, the FlowRepository identifier, or the ImmPort identifier of the dataset. The study ID reported in the first column can be used to locate the corresponding dataset in the supplementary data directory shared in this study. For example, the folder named "A001" in the supplemental data directory reports the data associated with the publication reported in the source column.

| Study ID | Source |
|---|---|
| A001 | https://pubmed.ncbi.nlm.nih.gov/39081632/ |
| A002 | https://pubmed.ncbi.nlm.nih.gov/32632085/ |
| A003 | https://pubmed.ncbi.nlm.nih.gov/32632085/ |
| A004 | https://pubmed.ncbi.nlm.nih.gov/25275464/ |
| A005 | https://pubmed.ncbi.nlm.nih.gov/32669297/ |
| A006 | https://pubmed.ncbi.nlm.nih.gov/32669297/ |
| A007 | https://pubmed.ncbi.nlm.nih.gov/32669297/ |
| A008 | https://pubmed.ncbi.nlm.nih.gov/26861911/ |

| | |
|---|---|
| A010 | https://pubmed.ncbi.nlm.nih.gov/33209300/#:~:text=Conclusion%3A%20As%20IL%2D10%2D,more%20severe%20COVID%2D19%20phenotype. |
| A011 | https://pubmed.ncbi.nlm.nih.gov/32632085/ |
| A012 | https://pubmed.ncbi.nlm.nih.gov/32632085/ |
| A013 | http://flowrepository.org/id/FR-FCM-Z2XF |
| A015 | https://pubmed.ncbi.nlm.nih.gov/39081632/ |
| A016 | https://pubmed.ncbi.nlm.nih.gov/30518691/ |
| A017 | https://pubmed.ncbi.nlm.nih.gov/26861911/ |
| A018 | https://pubmed.ncbi.nlm.nih.gov/26861911/ |
| A019 | Data shared by experiment author. |
| A020 | https://pubmed.ncbi.nlm.nih.gov/30862783/ |
| A021 | https://pubmed.ncbi.nlm.nih.gov/30862783/ |
| A022 | https://pubmed.ncbi.nlm.nih.gov/32830910/ |
| A030 | https://pubmed.ncbi.nlm.nih.gov/32717743/ |
| A032 | https://pubmed.ncbi.nlm.nih.gov/32807934/ |
| A039 | https://www.immport.org/ ID: SDY1743 |
| A041 | http://flowrepository.org/id/FR-FCM-Z27S |
| A044 | http://flowrepository.org/id/FR-FCM-Z36A |
| A045 | http://flowrepository.org/id/FR-FCM-Z2W3 |
| A046 | https://www.immport.org/ ID: SDY1389 |
| A048 | http://flowrepository.org/id/FR-FCM-Z2W4 |
| A049 | http://flowrepository.org/id/FR-FCM-Z27U |
| A053 | http://flowrepository.org/id/FR-FCM-Z2WZ |
| A054 | http://flowrepository.org/id/FR-FCM-Z2HV |
| A055 | http://flowrepository.org/id/FR-FCM-ZYXS |
| A056 | http://flowrepository.org/id/FR-FCM-Z2W6 |
| A058 | https://www.immport.org/ ID: SDY67 |
| A059 | http://flowrepository.org/id/FR-FCM-Z35D |
| A064 | https://pubmed.ncbi.nlm.nih.gov/33414483/ |
| A067 | https://pubmed.ncbi.nlm.nih.gov/32669287/ |

Supplementary Table 4: **Best cross-validation accuracy across all models tested.** The first column reports the name of the model tested: Random Forest (RF), feed forward neural network (FNN), K-nearest neighbor (KNN), decision tree (DT) and Naive Bayes (NB). The second column reports the best cross-validation accuracy observed. The last column reports the value of the parameters that gave the best cross-validation accuracy reported in the second column. In order, from top to bottom: Number of trees (n.tree) for the Random Forest, the number of neurons in the hidden layer (size) and the decay value (decay), the number of neighbours in the KNN (k), the value of the complexity parameter (cp) to determine tree length in the Decision Tree, the kernel density estimate status (activated or deactivated) for the Naive Bayes Model.

| **Model** | **Best accuracy** | **Best parameters** |
|---|---|---|
| RF | 89% | n.trees=2 |
| FNN | 47% | size=9,decay=1e-04 |
| KNN | 86% | k=5 |
| DT | 52% | cp=0.012 |
| NB | 52% | Kernel density estimate (True) |
